# Experimental and computational analyses of KEA3-dependent photosynthetic responses uncover species-specific molecular variation

**DOI:** 10.64898/2026.10.13.758443

**Authors:** Thekla von Bismarck, Quang Huy Nguyen, Jeremy Ruß, Linnéa Strandberg, Moritz Krämer, Julia Damen, Hans-Henning Kunz, Peter Jahns, Karin Köhl, Anna Matuszyńska, Ute Armbruster

## Abstract

- NPQ relaxation is thought to limit crop biomass and yield under natural, fluctuating light. The thylakoid K⁺/H⁺ exchanger KEA3 accelerates NPQ relaxation by lowering the lumen proton concentration and thus should affect biomass and yield under frequent light fluctuations.
- To investigate the impact of KEA3-dependent NPQ relaxation on *Solanum lycopersicum* (tomato), we generated knock-out and over-expression lines and measured NPQ, gas exchange, biomass and yield. We constructed a mathematical model to compare species-specific NPQ relaxation dynamics in tomato and the model plant *Arabidopsis thaliana*.
- In polytunnel and controlled fluctuating-light conditions, KEA3-mediated NPQ relaxation did not translate into consistent differences in tomato biomass or yield. Although KEA3 levels strongly affected NPQ relaxation, more so than in *A. thaliana*, the resulting effects on CO_2_ assimilation were minor. Modeling results indicated that, besides minor changes in KEA3 regulation, the activity of PsbS, the primary NPQ trigger, largely determines the species-specific differences in NPQ relaxation. Structural models from both species predict sequence variation with impact on PsbS activation.
- Manipulating KEA3 levels in tomato did not link NPQ relaxation with productivity. However, the results support a robust cross-species mathematical model that accurately predicts the molecular interactions governing NPQ dynamics.

## Introduction

Sunlight is the primary energy source for life on Earth, and its energy is captured through photosynthesis. Crop fields are planted densely to maximize yield per area, which creates a canopy where the upper leaves shade those below. Because sun angles shift and wind moves the foliage, this shading dynamically modulates light availability. Consequently, especially in dense stands, photosynthesis must frequently adjust to rapid and strong fluctuations in light, swiftly switching between light harvesting and photoprotection.

One major photoprotective mechanism of plants is the dissipation of excess absorbed light energy as heat, referred to as non-photochemical quenching (NPQ). This mechanism, which mainly takes place in the antenna of photosystem II (PSII (Nicol *et al*., 2019)), avoids the production of deleterious levels of reactive oxygen species (ROS, reviewed by (Demmig-Adams & Adams III, 2006; van Amerongen & Croce, 2025)). The most rapidly adjustable and main component of NPQ, the energy-dependent quenching qE, is activated by the acidification of the thylakoid lumen, which signals that absorbed light energy exceeds the requirements of downstream energy consuming processes. qE links light utilization to the proton motive force (PMF), which in chloroplasts is largely stored as a proton concentration difference (ΔpH) across the thylakoid membrane (reviewed by (Armbruster *et al*., 2017; Kunz *et al*., 2024)). Lumen pH is sensed by the PsbS protein via two glutamates, which both require protonation for full qE activation (Li *et al*., 2004). In addition, the lumen localized enzyme violaxanthin de-epoxidase (VDE) becomes active at low pH. VDE de-epoxidizes violaxanthin to form zeaxanthin that together with protonated PsbS activates the thermal dissipation process through rearrangements of the PSII supercomplex (reviewed by (van Amerongen & Croce, 2025)). Upon transition back to light limiting conditions, the lumen pH increases again, inactivating VDE while PsbS gets deprotonated, and the enzyme zeaxanthin epoxidase (ZEP) epoxidizes zeaxanthin to form the less qE active violaxanthin (reviewed by (Jahns & Bethmann, 2025)).

In *Arabidopsis thaliana* (*A. thaliana*), the thylakoid localized KEA3 K^+^/H^+^ antiporter was found to accelerate qE relaxation after transitions from high to low light by rapidly decreasing the proton concentration of the lumen without major effects on PMF (Armbruster *et al*., 2014, 2016). KEA3 activity is regulated via a stromal regulatory domain that responds to nucleotide ratios and stromal pH, thereby sensing the chloroplast energy state (Uflewski *et al*., 2024). Through this regulation, KEA3 activity is strongly affected by light intensity (Uflewski *et al*., 2021), but also by the light acclimation state of the plant (von Bismarck *et al*., 2023).

The processes that contribute to NPQ relaxation after transitions to light limiting conditions do not instantaneously shut down the dissipation of light energy as heat. It has been proposed that particularly in canopy settings, light energy for photosynthesis is lost due to lagging NPQ relaxation kinetics, thereby limiting crop yield (Zhu *et al*., 2004, 2008). Thus, improved NPQ relaxation rate is seen as potential pathway for photosynthesis and crop improvement.

One strategy to accelerate NPQ relaxation was the simultaneous overexpression of the two xanthophyll-cycle enzymes VDE and ZEP, together with PsbS (referred to as VPZ). VPZ improved plant biomass in tobacco and soy bean yield in the field (Kromdijk *et al*., 2016; De Souza *et al*., 2022), but it reduced biomass accumulation in *A. thaliana* (Garcia-Molina & Leister, 2020) and lowered the yield of the C_3_ crops potato and rice (Lehretz *et al*., 2022; Zheng *et al*., 2026). Xanthophylls share precursors with gibberellins and serve as intermediates of abscisic acid (ABA) biosynthesis, of which ZEP is a key enzyme (Koornneef *et al*., 1982; Niyogi *et al*., 1998; Park *et al*., 2008). Increased ABA levels were observed in the transgenic VPZ potato lines (Lehretz *et al*., 2022), supporting the hypothesis that VPZ can affect phytohormone pathways and thereby growth and reproduction (Kaiser *et al*., 2019). Therefore, species-specific differences in phytohormone levels and their effects may underlie the contrasting effects of VPZ across species.

The thylakoid K^+^/H^+^ antiporter KEA3 offers an alternative means to speed up NPQ relaxation (Armbruster *et al*., 2014, 2016). While KEA3 influences the de-epoxidation state of the xanthophyll pool under certain growth-light conditions, it does so by modulating lumen pH, which in turn affects VDE activity (von Bismarck *et al*., 2023). To date, no link has been established between lumen acidification and ABA or gibberellin levels, indicating that KEA3 changes NPQ relaxation without phytohormone-related side effects. In *A. thaliana*, *kea3* loss-of-function mutants displayed very slow NPQ relaxation kinetics, yet showed no measurable reduction in biomass under highly variable natural light, including polytunnel experiments (von Bismarck *et al*., 2023). This lack of biomass impact may reflect the species’ flat rosette architecture and fast-cycling ecology, which may limit the benefit of faster NPQ relaxation.

To test whether a more complex three-dimensional architecture alters the relationship between KEA3-mediated NPQ dynamics and productivity, we turned to tomato (*Solanum lycopersicum*), a fast-growing vegetable crop with a large, branched canopy. In tomato grown under polytunnel or controlled fluctuating-light conditions, KEA3-dependent NPQ relaxation did not translate into higher biomass or yield, mirroring the *A. thaliana* result, yet clear species-specific differences in NPQ relaxation and the effect of KEA3 were observed. We therefore combined comparative experiments with the mathematical modelling of NPQ dynamics during light fluctuations, allowing us to dissect the contributions of KEA3 and other NPQ components. Our analyses indicate that while strong sequence variation exists between the KEA3 proteins from tomato and *A. thaliana*, differences in the dynamics of PsbS protonation/ activation is the principal factor underlying the distinct KEA3-dependent NPQ responses of the two species.

## Material and Methods

### Identification and characterization of *SlKEA3*

The protein sequence of *A. thaliana* KEA3 (At4g04850) was used as a query to search for orthologs in *S. lycopersicum* using ITAG release 2.40 of the Sol Genomics Network. The proteins encoded by *Solyc11g044260.1* and *Solyc11g044250.1* showed significant sequence matches for the first 624 amino acids. Re-sequencing revealed that an annotated stop codon between *Solyc11g044260.1* and *Solyc11g044250.1* was not present and that both genes form a single reading frame. The sequence downstream of Solyc11g044250.1 was not annotated, and the 3’end of the *SlKEA3* cDNA was identified via 3’RACE with the SMARTer RACE 5’/3’ Kit (Takara Bio) according to the manufacturer’s instructions (see Table S1 for the gene-specific primer). The identified sequence of *SlKEA3* aligns with NCBI LOC101265888 (Buoso *et al*., 2022).

### Generation of *SlKEA3* knock-out and overexpression lines

To generate the *pTB001* vector for transgenic tomato selection using kanamycin as a selection marker, the *pEG001* (Chung *et al*., 2023) carrying a T-DNA with CRISPR/Cas9 cassette was modified by insertion of a kanamycin instead of the hygromycin resistance cassette between two *EcoR*I sites. Two pairs of *SlKEA3*-specific guide RNAs (gRNAs) were generated to target two different regions spanning 753 (pair 1) or 505 (pair 2) base pairs in the *SlKEA3* gene (Fig. 1a, Table S1). For each construct, a single fragment containing the gRNA scaffolding, U6 promoter, the specific pair of gRNA sequences, and U6 terminator (amplified from *pJF1046* (Ruf *et al*., 2019)) was inserted into *pTB001* as described in (Gonzalez-Duran *et al*., 2025). These constructs did not lead to the deletion of the entire targeted genomic regions, but to deletions of 6 bp at the binding site for the reverse primer of pair 1 in exon 11 (*slkea3-1*) and 1 bp at the binding site for the forward primer of pair 2 in exon 12 (*slkea3-2*). For the generation of mesophyll-specific *SlKEA3* overexpression lines, the promoter of the *Solanum tuberosum* cytosolic fructose-1,6-bisphosphatase (*StcyFBPaseP*) was used (Vallarino *et al*., 2020). A codon-optimized *StcyFBPaseP-SlKEA3cDNA-HA* sequence was synthesized by GENEWIZ (Germany) and inserted into the *Sbf*I and *Sal*I sites of the T-DNA sequence of the binary vector *pGWB2* upstream of the NOS terminator (Nakagawa *et al*., 2007). All constructs were validated by sequencing and then transformed into *Solanum lycopersicum* cv Moneymaker using *Agrobacterium tumefaciens* GV2260. Transformed tomato plants were selected based on their kanamycin resistance. *SlKEA3* overexpressors were selected based on the detection of the C-terminal HA epitope (YPYDVPDYA) using a specific HA-antibody at the expected molecular size of tagged SlKEA3 by immunoblotting. The CRISPR/Cas9 generated *slkea3* knock-out lines were identified by PCR amplification over the gRNA target sites (as described in (Bhattacharya & Van Meir, 2019)) and sequencing of the product.

**Figure 1.**
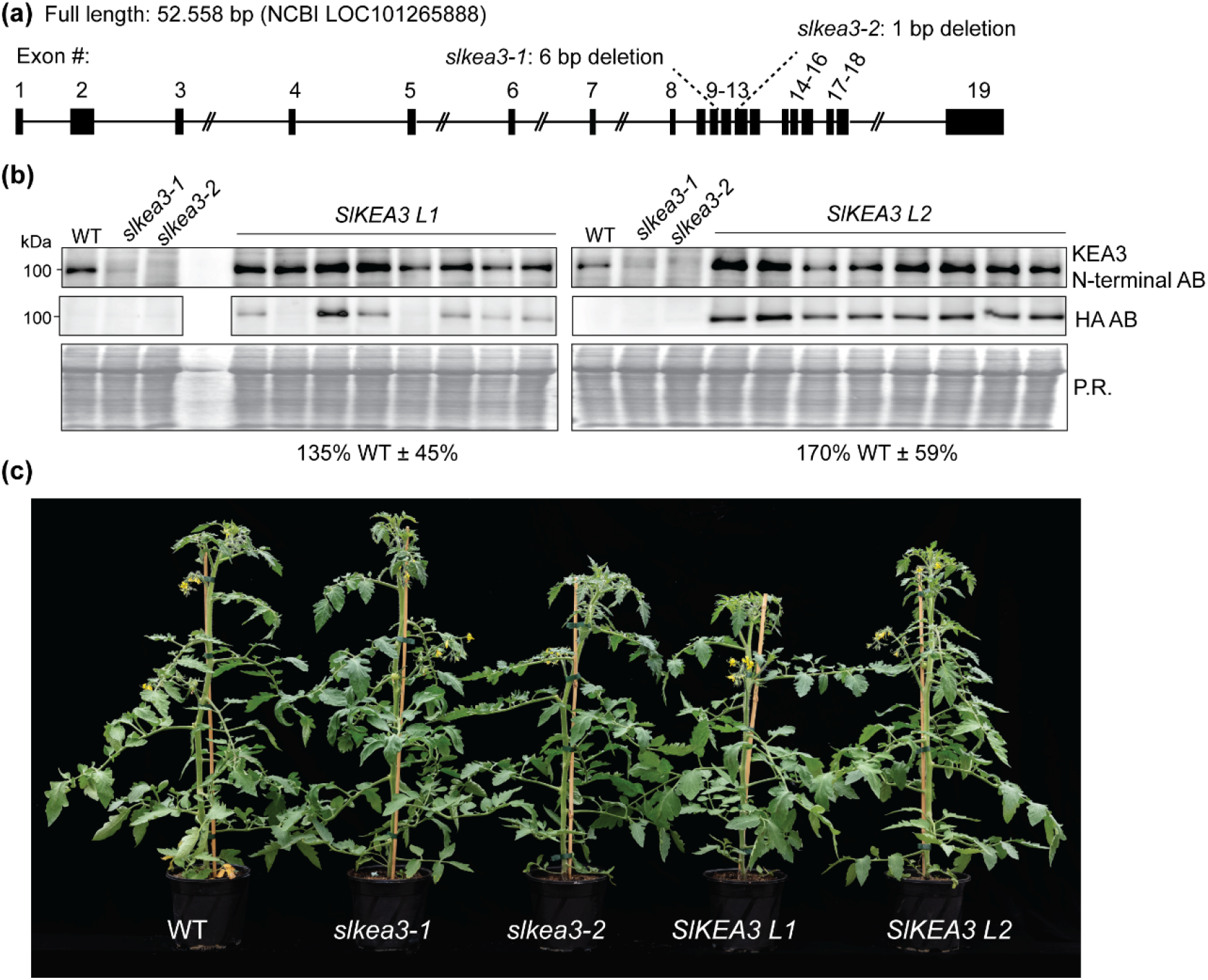
Generation of *SlKEA3* knock-out and overexpression lines. (a) Gene map of the *SlKEA3* gene based on own sequence analysis and 3’ mapping aligning with NCBI locus 1012655888, including information on *slkea3-1* and *slkea3-2* mutants generated by Crispr/Cas9. *slkea3-1* harbours a 6 bp deletion at the guide-RNA target site of sgRNA-rev1, leading to a deletion of a threonine and glutamine within transmembrane helix 6 of SlKEA3 at position 367 and 368. *slkea3-2* had a 1 bp deletion at the target site of the sgRNA-fwd2 guide-RNA, resulting in a frameshift and a premature stop codon. (b) Immunoblot analysis of total leaf protein extract from Money Maker WT and *SlKEA3* knockout (*slkea3-1* and *slkea3-2*) and overexpression (*SlKEA3 L1* and *SlKEA3 L2*) lines using antibodies against an N-terminal AtKEA3 epitope sequence conserved in SlKEA3 and the HA-tag of SlKEA3-HA in the overexpression lines. Ponceau red (P.R.) staining of the membrane after transfer is shown as an experimental control and was used for normalization of SlKEA3 levels for the quantification relative to WT-levels shown below the images (Average ± SD in %). KEA3 quantification for the overexpression lines was only included, when the HA immunoblot gave a positive result (c) Picture of representative 8-week-old tomato plants from each genotype grown in the polytunnel from 09.08.2022 to 07.10.2022 (PT1).

### Plant material and Growth conditions

Tomato plants were grown on peat-based substrate in 13 cm pots, except for polytunnel experiments, in which plants were pre-cultivated in 6 cm pots for 25 d (PT1) and 27 d (PT2) in a phytochamber (16 h of approximately 120 μmol photons m^−2^ s^−1^) and then transferred to 18 cm pots for growth in the polytunnel from 09.08.2022 to 04.11.2022 (PT1) and 11.05.2023 to 18.07.2023 (PT2). Plants were fertilized with 5 g Entec perfect (14-7-17) on potting and 4 g Entec perfect again at the onset of flowering. Environmental data, i.e. light intensity, relative air humidity and air temperature was logged (see Table S2). For the random fluctuating (RL) experiments with a 16 h light period, a climate chamber was set to 20°C and 60 % in the light and 16°C and 75 % in the dark. The RL regime was generated as described in von Bismarck *et al*., 2023 using white LEDs with an average light intensity of 250 μmol photons m^−2^ s^−1^. All plants were watered every 2-3 days. Drought-treated plants received water only once per week, except for the ionomics experiment, during which drought stressed plants were watered with half the volume of the control plants starting 6 weeks after germination. *A. thaliana* Col-0 and *kea3-1* and *kea3-2* (von Bismarck *et al*., 2023) were cultivated in 6 cm pots under the same RL conditions, except with a 12 h/ 12 h day/ night cycle. The matching of DLI (day light integral) between *S. lycopersicum* and *A. thaliana* RL experiments at different day length led to differences in light intensity.

### Biomass and yield quantifications

In all conditions, the side shoots were removed upon appearance. Plant biomass was determined when the flowers of the first inflorescence of most plants had opened. To this end, leaves and stems were cut separately and dried at 65 °C for a minimum of 7 days. To determine the fruit yield of PT plants, the shoot tip was removed above the 2^nd^ inflorescence 11 weeks after sowing (PT1) or above the 3^rd^ inflorescence 12 weeks after sowing (PT2). The number and weight of all tomato fruits per plant was determined 6 (PT1) or 2 weeks (PT2) after removing the shoot tip.

### Spectroscopic measurements

An Imaging-PAM Maxi (Walz GmbH, Effeltrich, Germany) was used for chlorophyll (Chl) *a* fluorescence measurements during a fluctuating light set-up in a climate-controlled darkened spectroscopy room. All measurements were done on the 2^nd^ or 3^rd^ leaflets of the 3^rd^ or 4^th^ compound leaf (counting from the top all leaves longer than 5 cm), unless stated otherwise. Whole plants were put in the dark for 30 min before attached leaves were measured using the Imaging PAM (RL, PT2). For PT, leaflets were cut off after the dark period and placed in a wet petri dish before determining the minimal fluorescence in the dark (F_0_) and the stationary fluorescence (F_s_) in the light. Saturating pulses (SP, setting 10) were used to determine the maximum fluorescence in the dark (F_m_) and in the light (F_m_’). Fluctuating light regimes were applied, alternating between 4 min low light (200 µmol photons m^-2^ s^-1^) and 1 min high light (2000 µmol photons m^-2^ s^-1^), and a final 5 min dark period. The SPs were applied every 20-40 seconds. NPQ was calculated as (F_m_- F_m_’)/ F_m_’ and Φ_PSII_ as (F_m_’-F_s_)/F_m_’.

MultispeQ devices (Kuhlgert *et al*., 2016) were used for measurements of Chl *a* fluorescence and absorbance, using the Photosynthesis Rides 2.0 program (https://openjii.org/en-US/platform/experiments/e0acb807-5a1a-4f35-8902-a1d8629d4cc9/dashboards/461fd3d0-0844-471a-8ddc-2f490000d3f0). ECS_t_ was re-calculated in Python by fitting a single exponential decay to the slope of the 520 nm absorbance change after the transition from light to dark using scipy.optimized. Detailed description of experiment and data processing can be found on the OpenJII dashboard (link above). For the RL drought experiments, measurements were conducted before watering and after one and two weeks of drought stress. All *in vivo* measurements were done in a randomized fashion, starting earliest 2 h after the light was switched on/dawn and latest 2 h before the light was switched off/dusk.

### Gas exchange measurements

The GFS-3000 open gas exchange system with the LED array unit 3056-FL (with 90% red and 10% blue light) connected to the optical oxygen sensor 3085-O2 (Walz GmbH, Effeltrich, Germany; Flow: 750 µmol s^−1^, CO_2_ concentration: 400 µmol, relative humidity: 60%, cuvette temperature: 20 °C) were used for gas exchange measurements in 2 s intervals on the 2^nd^ or 3^rd^ leaflet of the 4^th^ leaf (see above). Leaf size, determined from photos via ImageJ, was used to normalize the data. The same light regime as for the Chl *a* fluorescence measurements was used but extended to 45 min to assure reaching of a pseudo steady state during the light fluctuations. Due to watering problems in the controlled climate chamber, single plants might have not been watered sufficiently which manifested in a low stomatal conductance (*g_s_*) that led to very low CO_2_ assimilation. Data was excluded when *g_s_* dropped below 200 mmol m^-2^ s^-1^ for RL. This cut off was chosen by comparing to the average of *g_s_* in this condition.

### Bioinformatics and Statistics

Sequence homology searches were performed using NCBI BLAST (blastp) with the SlKEA3 amino acid sequence as a query. Searches were performed against the NCBI Protein Reference Sequences (refseq_protein, update date: 29.04.2026) database with default parameters. The first 15 best hits were used and redundant sequences were excluded. Sequence alignments were visualized using the NCBI Constraint-based Multiple Alignment Tool (Cobalt) with colouring based on conservation with default settings (conservation setting: 2 bits). The models of KEA3 dimers and PsbS were predicted by the AlphaFold server (https://alphafoldserver.com; (Abramson *et al*., 2024)) with version AlphaFold 3, and molecular graphics were performed with UCSF ChimeraX v1.11.1 (Pettersen *et al*., 2004). Statistical analyses were performed in Python (v 3.13.9) using the SciPy and statsmodels packages.

### Protein extraction and immunoblotting

For total protein extraction, ground leaf material (leaf 4-5) was mixed with extraction buffer (200 mM Tris, pH 6.8, 8% (w/v) SDS, 40 % (v/v) glycerol, and 200 mM DTT) to a concentration of 0.1 mg/ µl buffer) and heated at 65 °C for 10 min. Proteins were separated by size via SDS-PAGE and blotted onto nitrocellulose. Protein loading was visualized with Ponceau Red (0.1% (w/v) Ponceau S in 5% (v/v) acetic acid). For immunodetection of AtKEA3 and SlKEA3, the N-terminal antibody against AtKEA3 as described in (Uflewski *et al*., 2021) against the epitope SAVDVINDLGFDTLT was used. This 15 amino acid sequence is conserved in SlKEA3 except for an L instead of an F at position 11. The antibody against the HA-epitope was from Agrisera (AS122220). SlKEA3 levels were normalized to wild type levels and the ponceau red stain of the membrane using ImageJ.

### Quantification of leaf element contents

For element analyses, leaf tissue was dried at 67°C for a minimum of 7 days. Sample preparation was carried out according to (Holzner *et al*., 2026). In brief, dried leaf material was homogenized using a zirconium mortar (Stanford Advanced Materials, Santa Ana, CA, USA). Using an ultra-balance (Sartorius, Göttingen, Germany), aliquots of 3 mg were taken and subsequently digested in 200 ul of 69% (v/v) ultra-pure HNO_3_ (Roth, Karlsruhe, Germany) at 95°C for 90 min. Digested samples were diluted 1:10 with element standards to a final concentration of 50 ppm Scandium (WL) and 1 ppm Gallium (Mo-K) (Sigma Aldrich/Merck, Darmstadt, Germany). All samples were spotted on silicon-coated quartz glass carriers (B&M Optik, Limburg, Germany) and measured by total reflected X-ray fluorescence (TXRF) on a S4 T-STAR (Bruker, Berlin, Germany) at 50 kV, for 600 s per sample using Mo-K and W-L excitation. Subsequently, the element amount per sample was normalized to dry weight.

### Pigment extraction and quantification

Leaflet 2 and 3 of leaf 4 of the tomato plants and the whole *A. thaliana* rosettes were harvested in the middle of the light period and immediately frozen in liquid nitrogen. For RL experiments, the light intensity was set to 300 µmol photons m^-2^ s^-1^ during harvest. Whole leaf or whole rosette pigments were extracted from 20 mg ground material in the dark and on ice by two times adding 500 µl 100% acetone, mixing well and centrifuging at 14,000 rpm for 10 min at 4°C. The pooled supernatants from both extraction rounds were filtered through 0.2 µm Sartorius Minisart SRP15 PTFE filter. Pigment extracts were quantified via HPLC as described in (Färber *et al*., 1997).

### Mathematical, kinetic model of NPQ and numerical simulations

A kinetic model based on ordinary differential equations (ODEs) was developed to describe NPQ and the light reaction. The model expands on earlier NPQ formulations (Matuszyńska *et al*., 2016), with adjustments based on (Cruz *et al*., 2001; Li *et al*., 2021) coupling PSII and NPQ kinetics with PMF partitioning, and KEA3-dependent K^+^/H^+^ exchange. The model thereby links Chl *a* fluorescence-derived measurements to the underlying lumen pH, membrane potential, PsbS protonation state, and KEA3 transport activity. The model includes a simplified photosynthetic electron transport chain comprising PSII, cytochrome *b*_6_*f*, ATP synthase, and KEA3, with NPQ represented through the PsbS- and xanthophyll-dependent components. A complete description of the model is provided in Supplementary Material (Method S1). Most biochemical and physical parameters were adopted from literature or previous models (Method Table S1). A limited subset of species-dependent parameters was adjusted to experimental data and used as reference parameter sets, then the residual was calculated using a multi-objective residual function that consider deviations in NPQ and Φ_PSII_ kinetics across genotypes, accounting for species-specific responses as well as differences between genotypes (Table S3). To assess model robustness, ∼ 10,000 random parameter sets were sampled by Monte Carlo perturbation within a ±10% neighbourhood of the fitted solution, and their residual performance was compared to the reference set residual. This analysis tested if the fitted parameter sets represented locally robust solutions.

## Results

### Generation of *SlKEA3* knock-out and overexpression lines

We generated *slkea3* CRISPR/Cas9 knock-out and *SlKEA3* overexpression lines in the variety Money-Maker (M.M.). Two *slkea3* lines deriving from independent sets of guide-RNA pairs were selected (Fig. 1a, Table S1). Immunoblot analysis using the N-terminal epitope AtKEA3 antibody (Uflewski *et al*., 2021) revealed a signal at the predicted molecular weight of 100 kDa in the M.M. WT, of which levels were strongly decreased in *slkea3-1* and absent in *slkea3-2* (Fig. 1b). The overexpression lines were generated by fusing *SlKEA3* cDNA with a C-terminal HA-tag to the mesophyll-specific *StcyFBPase* promotor. Two independent lines, *SlKEA3 L1* and *L2* were selected based on the presence of an HA-signal at the predicted molecular weight of 105 kDa. In the next generation, SlKEA3 was quantified via immunoblot analysis (Fig. S1, Table S4). Both *SlKEA3 L1* and *L2* only showed minor increases in SlKEA3 compared to M.M., with an average of 135% and 172%, respectively.

The *SlKEA3* knock-out and overexpression lines were grown together with the wildtype M.M. in two independent polytunnel (PT) trials under natural fluctuating light and climate conditions resulting in similar plant growth phenotypes (Fig 1c, see Table S2 and Fig. S2 for logged climate data during PT1 and PT2).

### The dynamics of NPQ are dependent on SlKEA3 levels and growth light

Next, we measured Chl *a* fluorescence during strong light fluctuations to determine NPQ (Fig. 2a-b, Fig. S3-5) and the quantum yield of photosystem II (Φ_PSII,_ Fig. 2c-d, Fig. S3-5). At first glance, tomato *KEA3* lines showed a similar phenotype to previously published *A. thaliana* mutant and overexpressors (Armbruster *et al*., 2014, 2016): *slkea3* had much higher NPQ after transition from high to low light, while *SlKEA3* overexpression lines had slightly lower NPQ (Fig. 2a-b). The increase in NPQ in the *slkea3* knock-out lines was accompanied by a decrease in Φ_PSII_, while only in one of two polytunnel experiments (PT1) the average decrease in NPQ in the *SlKEA3* lines was accompanied by an average increase in Φ_PSII_ (Fig. 2c, d). In PT2, Φ_PSII_ of the two *SlKEA3* lines was comparable to M.M (see Fig. S4-5 for more detailed analyses). These results suggested that the dependence of Φ_PSII_ on NPQ in *SlKEA3* lines is affected by the specific environmental conditions during growth. These were very different between the two trials: PT1 was sown late in the season and light and temperature was moderate, while there was a heat wave and temporary water stress during the PT2 trial (Fig. S2, Table S2).

**Figure 2.**
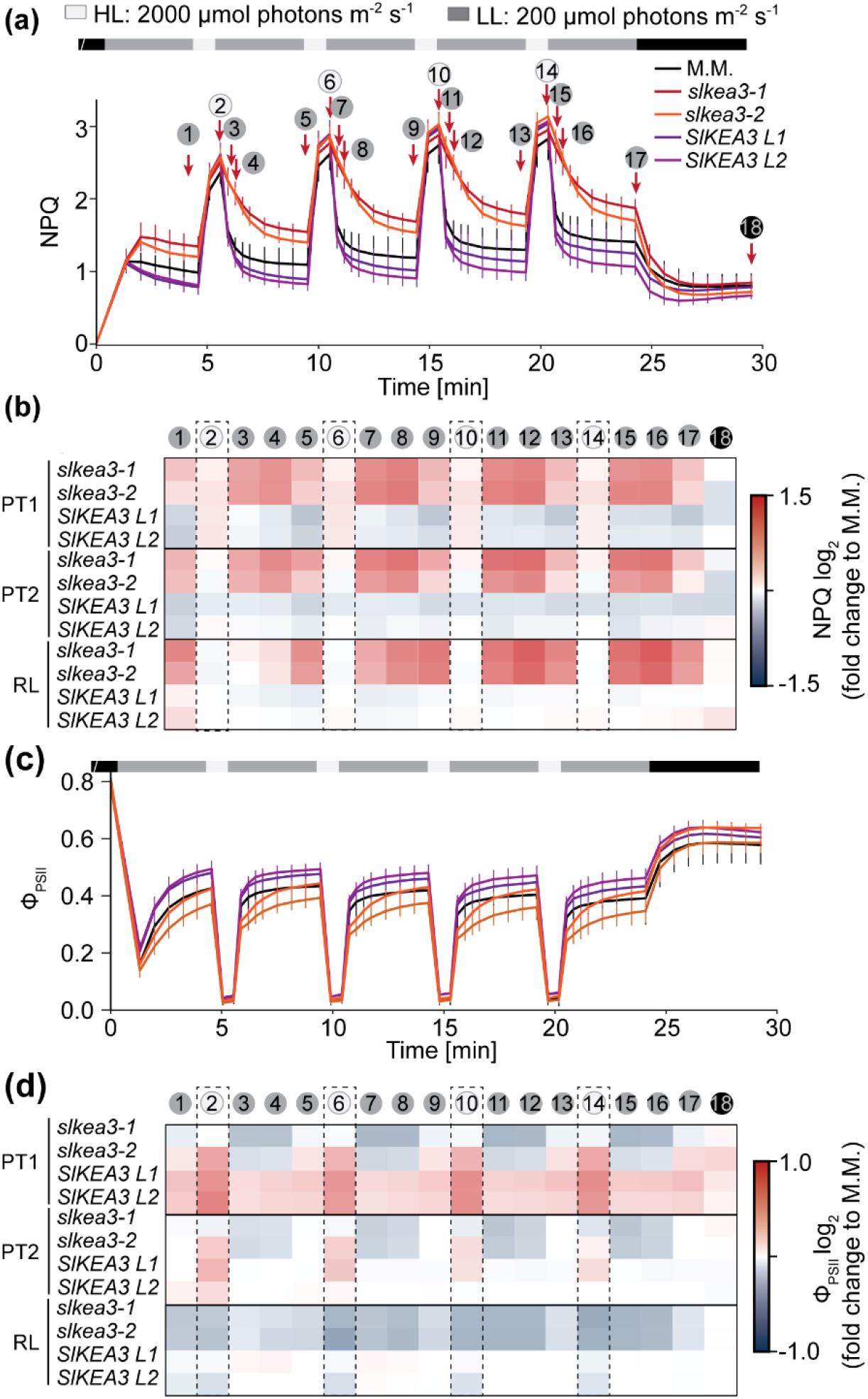
Analysis of Chl *a* fluorescence derived parameters during light fluctuations. (a, c) Traces of non-photochemical quenching (NPQ; a) and photosystem II quantum efficiency (Φ_PSII_; c) of 11-week old wildtype Money Maker (M.M.), *SlKEA3* knock-out and overexpression lines grown in the polytunnel (PT1). Shown are averages ± SD of N=5 (*SlKEA3 L1*) or N=7 (WT, *slkea3-1*, *slkea3-2*, *SlKEA3 L2*). (b, d) Numbers in circles in (a) indicate timepoints used for heatmap representation of log_2_ transformed NPQ (b) and Φ_PSII_ (d) of plants from PT1, 9-week-old plants from PT2 and 7-week-old plants from random fluctuating light (RL). Colouring is based on log_2_ transformed average ratios to M.M. with N=4 (RL: *slkea3-1*), N=5 (PT2: *SlKEA3 L1*, *SlKEA3 L2*; RL: WT, slkea3-1, *SLKEA3 L2*), N=6 (RL: *SlKEA3 L1*), N=7 (PT1: WT, *slkea3-1*, *slkea3-2*, *SlKEA3 L2*, PT2: WT, *slkea3-1*, *slkea3-2*). For PT1 and PT2, detached leaves were used for measurements and attached leaves for RL. Phenotypes were similar between attached and detached leaves, as shown in Fig. S3 and S4 for PT2.

To control for the climate, we grew the tomato lines in a growth chamber, set to daily changing, randomly fluctuating light (RL) with a constant daily light integral. The *slkea3* plants showed very slow NPQ relaxation compared with M.M., accompanied by decreased Φ_PSII_, but no differences could be observed for *SlKEA3* (Fig. 2, Fig. S3).

### Slow NPQ relaxation only poses a weak penalty on CO_2_ assimilation

To assess the effect of slow NPQ relaxation and Φ_PSII_ recovery on net CO_2_ assimilation (A_net_), we conducted gas exchange measurements during light fluctuations (Fig. 3a-b, Fig. S6). Both PT1- and RL-grown *slkea3* plants showed a transient reduction in A_net_ relative to M.M. after the high-to-low light transition. The two *SlKEA3* lines from both conditions behaved similarly to their respective M.M., with average differences falling within experimental noise (Fig. S6). Although *slkea3* mutants exhibited ∼2 times the NPQ of M.M. throughout the low-light phases of RL, the accompanying transient decrease in A_net_ never exceeded ∼25 % (Fig. S7). Thus, only a small fraction of the difference in NPQ was reflected in CO_2_ assimilation.

**Figure 3.**
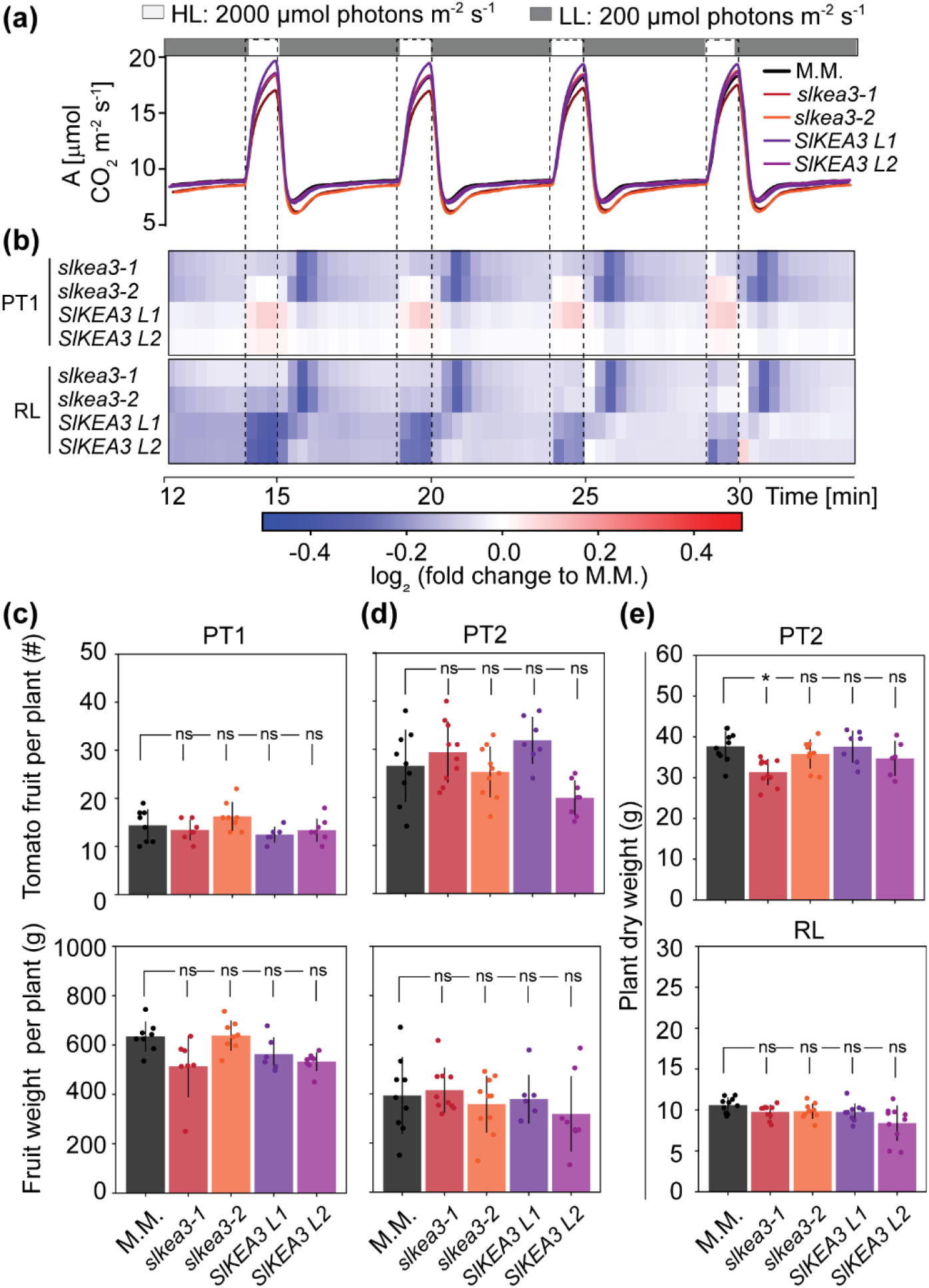
The differences in carbon fixation rates do not translate into changes in yield or growth. (a) CO_2_ assimilation rates (A_net_) of 11-week old wildtype Money Maker (M.M.), *SlKEA3* knock-out and overexpression lines grown in PT1 were measured and light fluctuations 3-6 are shown. Full traces can be found in Fig. S6. (b) Heatmap representation of log_2_ transformed A_net_ changes of PT1 plants shown in (a), and 7-week-old plants from random fluctuating light (RL). CO_2_ assimilation rates were binned over 20 seconds. Colouring is based on log_2_ transformed average ratios to M.M. with N=5 (PT1: *SlKEA3 L1*, RL: SlKEA3 L1, *SlKEA3 L2*), N=6 (RL: *slkea3-1*, *slkea3-2*) and N= 8 (PT1: WT, *slkea3-1*, *slkea3-2*, *SlKEA3 L2*; RL: WT). (c-d) Tomato yield was determined from the PT1 experiment (c) of 12-week-old plants and PT2 of 8-week old plants (d) as number of fruits per plant and total fruit weight per plant. Bars represent mean ±SD of N=6 (PT1: SlKEA3 L1, PT2: WT), N=7 (PT1: *slkea3-1*), N=7 (PT2: *SlKEA3 L1*) N=8 (PT1: WT, *slkea3-2*, *SlKEA3 L2*; PT2: *SlKEA3 L2*), N=10 (PT2: WT, *slkea3-1*, *slkea3-2*) (e) Total plant dry weight of 9-week-old plants from PT2 and 8-week-old plants from RL. Bars represent mean ± SD of N=6 (PT2: *SlKEA3 L1*, *SlKEA3 L2*), N=7 (RL: *SlKEA3 L2*), N=8 (RL: *SlKEA3 L1*), N=10 (PT2: WT, *slkea3-1*, *slkea3-2*; RL: WT, *slkea3-1*, *slkea3-2*). (c-e) Data were analyzed using one-way ANOVA followed by Tukey’s post hoc test. When data did not meet normality and homogeneity of variances criteria for ANOVA, a Kruskal–Wallis test followed by Dunn’s post hoc test was applied. Asterisks indicate significant differences to M.M.. ns, not significant.

While the differences in A_net_ during a single light fluctuation were modest between *slkea3* and M.M., repeated reductions in A_net_ throughout the day and across the growing season, particularly under the highly dynamic light conditions of RL and the PT trials, could accumulate and ultimately limit biomass production and potentially yield. However, no consistent difference in tomato yield (Fig. 3c, d) or plant dry weight (Fig. 3e) could be observed between *slkea3* and M.M.

### SlKEA3 is deactivated in drought stress conditions

Under drought stress, stomata close to reduce further water loss, decreasing CO_2_ availability and CBB cycle activity. *A. thaliana kea3* mutants were reported to show stronger growth deficits than WT under drought (Yamada *et al*., 2024). To investigate this potential link between KEA3 function and drought resistance in this agronomically relevant vegetable crop, we performed drought stress experiments on 4-week-old tomato WT and *slkea3* (Fig. 4a-b). Simultaneous measurements of NPQt, which approximates total NPQ without initial dark acclimation phase, particularly useful for field experiments (Tietz *et al*., 2017), and ECSt, which serves as a proxy for light induced PMF (Takizawa *et al*., 2007), were carried out directly in the chamber. Well-watered *slkea3* had in general higher NPQt values per ECSt than M.M., suggesting that KEA3 was active in the latter, decreasing the proton concentration in the lumen and thereby NPQt (Fig. 4 c-d). These results mirror data from *kea3* mutants in *A. thaliana*, which were further validated by spectroscopic PMF partitioning measurements (Armbruster *et al*., 2014; von Bismarck *et al*., 2023). In drought treated plants however, the difference in NPQt per ECSt was no longer present between *slkea3* and M.M., suggesting that KEA3 was inactive in M.M.. These results from drought treatments reinforce the notion that KEA3 is only active under conditions, in which both ATP and NADPH are efficiently turned over by the CBB cycle to increase light utilization (von Bismarck *et al*., 2023; Uflewski *et al*., 2024). While plant growth was severely decreased by the drought treatment, again no difference in plant size and dry weight was observed between *slkea3* and M.M. (Fig. 4e-f).

**Figure 4.**
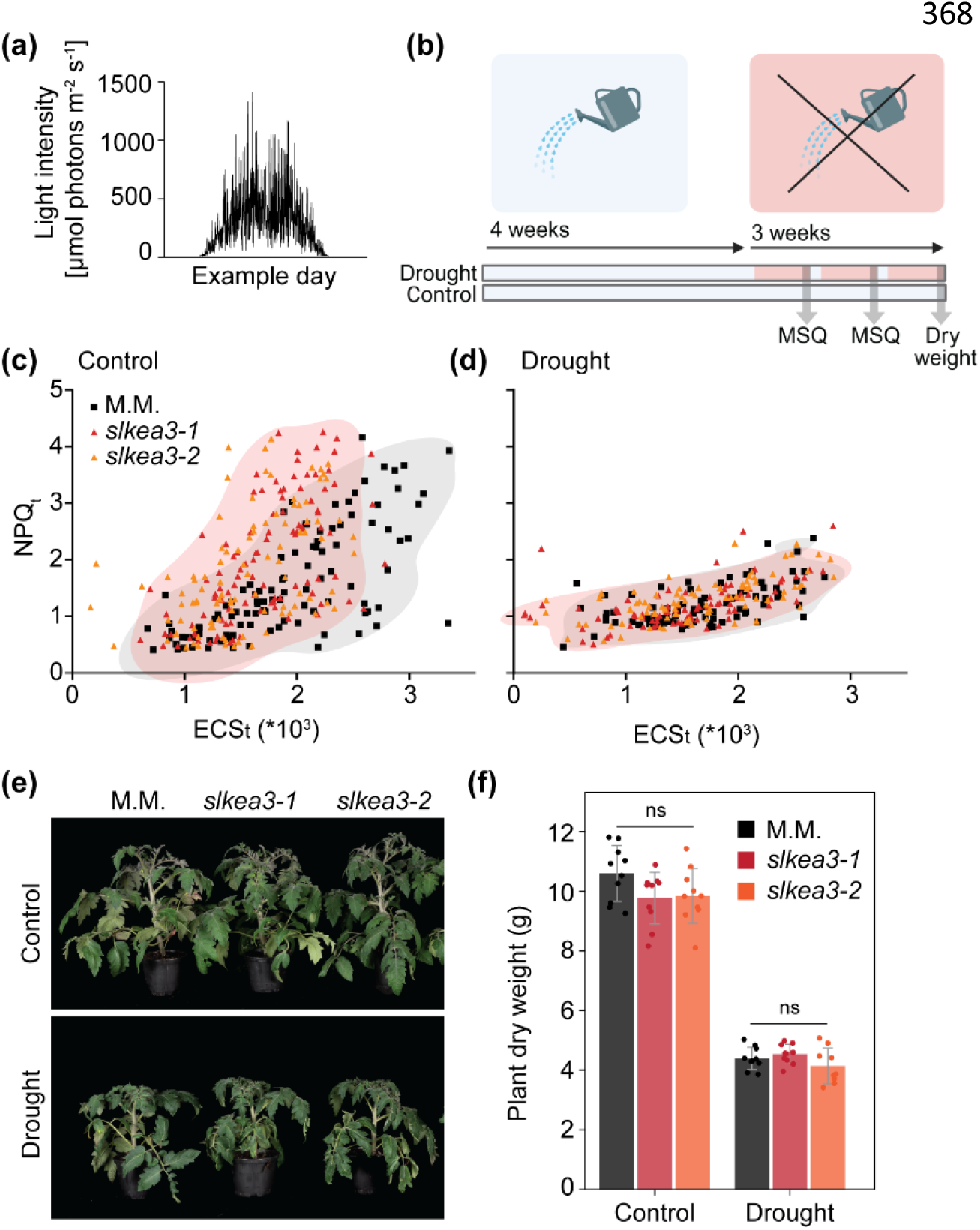
During drought stress, tomato KEA3 is switched off. (a) Example day of the light regime in random fluctuating light (RL). (b) Sketch of the drought stress treatment. Both control and drought stressed plants were watered every 2-3 days for the first 4 weeks after sowing. Drought stressed plants were then only watered every 7^th^ day for three weeks. (c-d) Steady-state NPQ_t_ versus ECSt measured after 5 and 6 weeks of control (c) and drought-stressed plants before watering (d). NPQ_t_ and ECSt data was filtered to include only data between 0-6 and 0-0.004, respectively. Kernel density estimates (KDE) with a density of >10% of max density are overlaid to indicate the density of points: grey for M.M and red for both *slkea3-1* and *slkea3-2*. Plots are based on N=97-106 (Control) and N=86-87 (Drought). (e) Pictures of 7-week-old representative WT (Moneymaker, M.M.) and *slkea3* plants. (f) Dry weight of plants as in (e). Shown is the average of N=10 ± SD. Data from control conditions are identical to M.M., *slkea3-1* and *slkea3-2* shown in Fig. 3e. Data were normally distributed and showed equal variance and thus were analyzed using one-way ANOVA followed by Tukey’s post hoc test, revealing no significant (ns) differences between genotypes.

Also the plant ionome was not affected by loss of KEA3, while treatment- and leaf-age dependent effects were found (Fig. S8-9).

### SlKEA3 differs from AtKEA3 by a C-terminal extension

By mapping the *SlKEA3* sequence, we found that the coding sequence was extended at the 3’ end when compared to *AtKEA3*, adding 200 amino acids to the C-terminus (Fig. 5a). Sequence analysis of closely related species showed that this C-terminal extension can be found in all analyzed *Solanum* species (Fig. S10), including *Solanum tuberosum* (potato). This sequence addition was confirmed via immunoblot analysis demonstrating that the molecular weight of SlKEA3 is ∼30 kDa higher than AtKEA3 (Fig. 5b).

**Figure 5.**
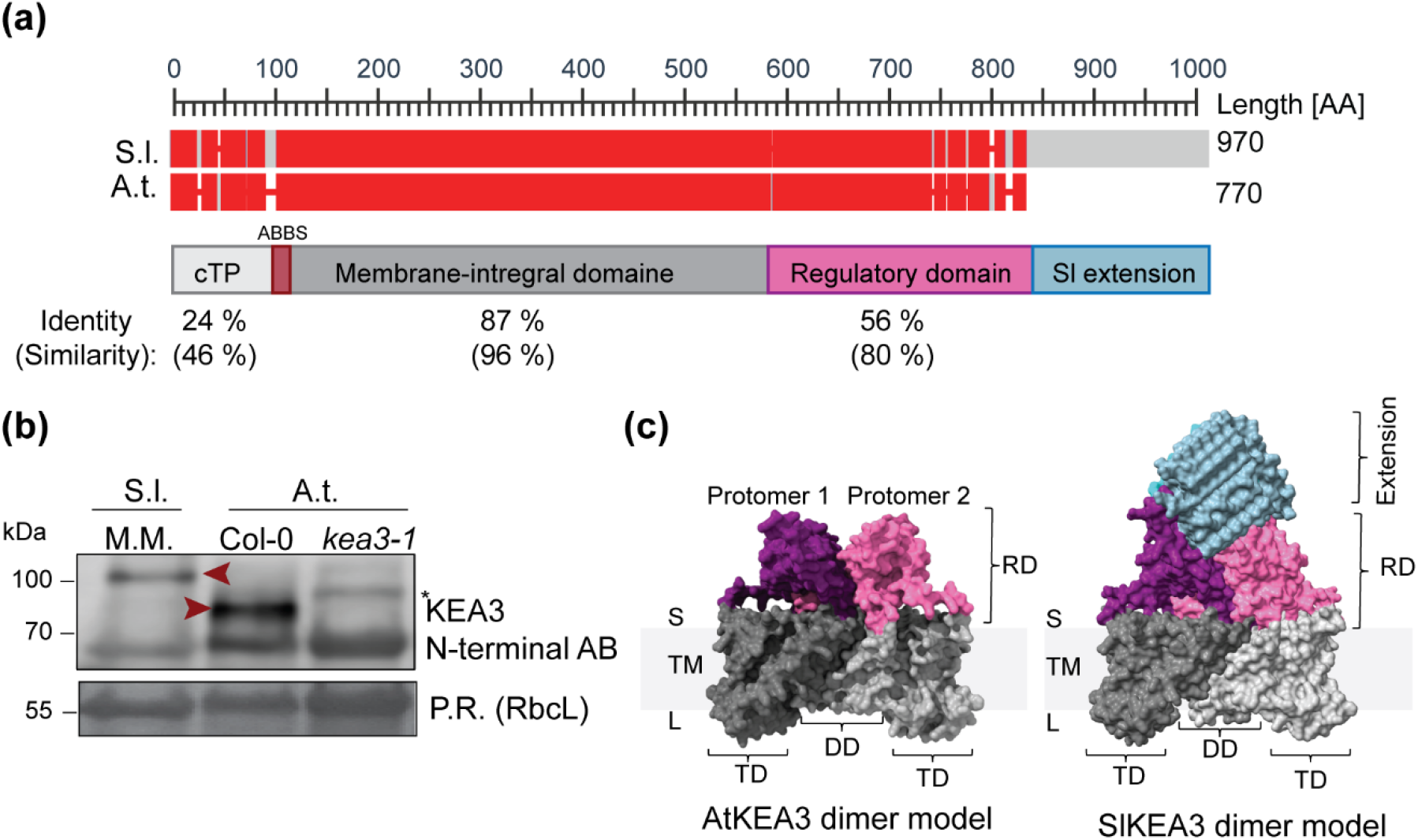
Comparison of KEA3 protein sequence and structure between *S. lycopersicum* and *A. thaliana.* (a) The KEA3 protein from *S. lycopersicum* (S.l.) and *A. thaliana* (A.t.) are conserved, but SlKEA3 but has an additional C-terminal extension (Sl extension). AA – Amino acid, cTP – Chloroplast transit peptide, ABBS- Antibody binding site. (b) Immunoblot analysis of total leaf protein extract from S.l. Moneymaker WT and A.t. Colombia-0 (Col-0) WT and A.t. KEA3 knockout mutant *kea3-1* using the N-terminal antibody (AB) as in Fig. 1B confirms proteins differ in size by ∼30 kDa. Ponceau red (P.R.) staining of the membrane after transfer is shown as an experimental loading control. (c) AlphaFold3 modelling of two protomers predicts both proteins form dimers with the Sl extension extruding at the stromal side. S – Stroma, TM – Thylakoid membrane, L – Lumen, RD – Regulatory domain, TD – Transmembrane domain, DD - Dimerization domain.

AlphaFold3 predicted both SlKEA3 and AtKEA3 to form dimers with a main interface of the two protomers in the transmembrane region (Fig. 5c), in agreement with a recent cryo-EM structure of the KEA3 homolog KefC from *E. coli* (Gulati *et al*., 2024). Notably, the SlKEA3 C-terminal extension is highly repetitive and predicted to form an extensive β-sandwich with 7 β-sheets on each side (Fig. S11). It is joined to the C-terminus of the nucleotide-binding regulatory domain of KEA3 that faces the stromal side of the thylakoid membrane.

### Loss of KEA3 has conserved but distinct effects on NPQ dynamics in Arabidopsis and tomato

Next up, we compared *A. thaliana* and tomato WT and knock-out mutants grown under RL conditions. Overall WT NPQ was lower in tomato than in *A. thaliana* (Fig. 6a; Fig. S12). Notably, the *slkea3* mutants showed a more strongly elevated NPQ during low light periods compared to WT. Thus, KEA3 suppresses NPQ more strongly in tomato than *A. thaliana* when plants undergo a transition from high to low light. Given that SlKEA3 carries a large C-terminal stromal extension that AtKEA3 lacks, differences in transport activity or regulation may underlie the stronger KEA3 effect on low light NPQ in tomato.

**Figure 6.**
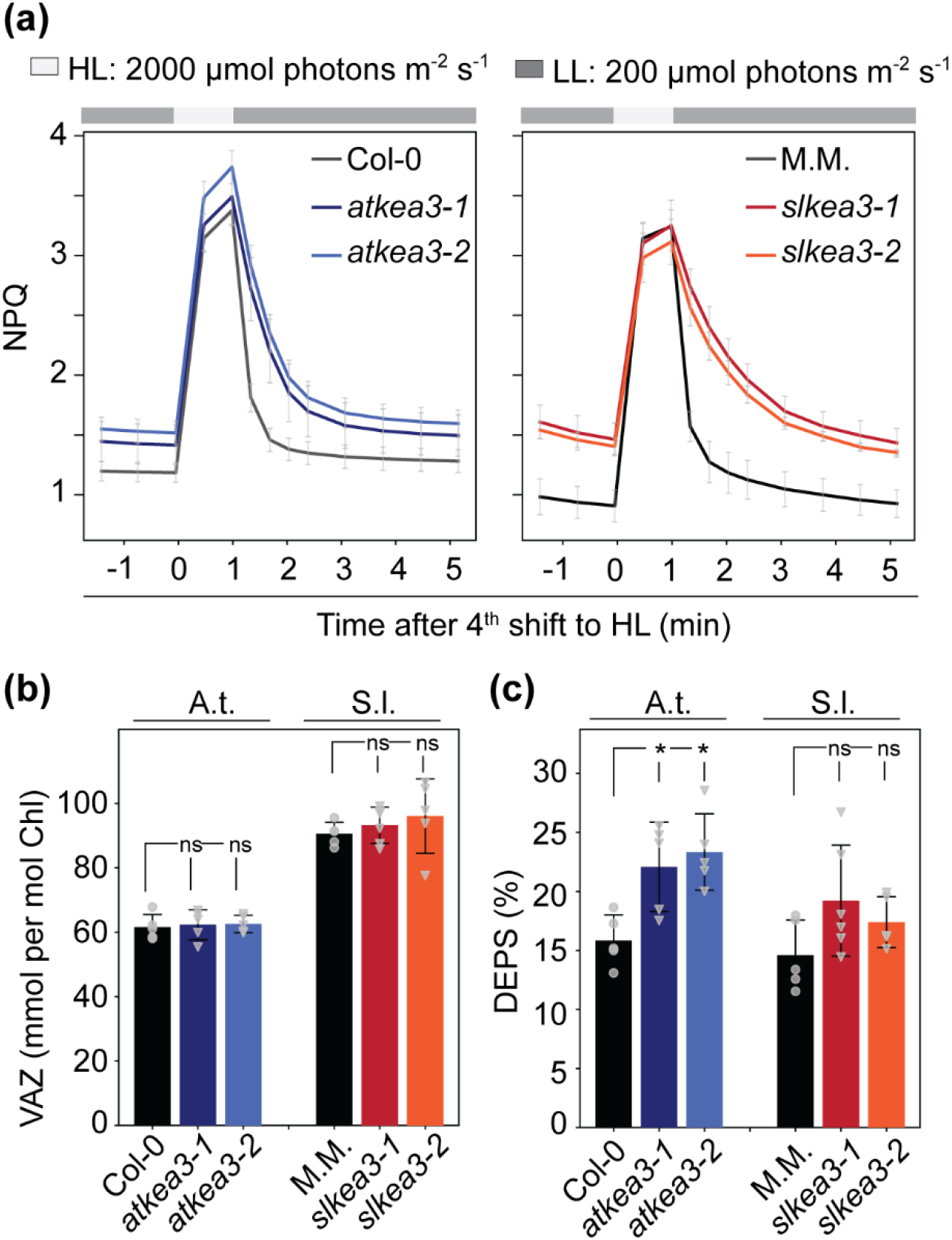
Comparative physiological analyses of *kea3* mutants in *A. thaliana* and *S. lycopersicum* reveal some differences. (a) Chlorophyll *a* fluorescence was measured on 30 min dark adapted whole rosettes of *A. thaliana* and 2^nd^ or 3^rd^ attached leaflet of leaf 3 or 4 of *S. lycoperscium* plants grown under RL with the same average daily light intensity of 250 µmol photons m^-2^ s^-1^ and species-adjusted daylength (12 h for A.t., 16 h for S.l.). The average NPQ during the 4^th^ light fluctuation is shown as mean ± SD for N=4 (S.l.: *slkea3-1*, *slkea3-2*) and N=6 (A.t.: all; S.l: M.M.). (b-c) Pigment analysis was performed on tissue samples, harvested in the middle of the light period, and the xanthophyll VAZ pool (violaxanthin + antheraxanthin + zeaxanthin, b) and its deepoxidation state (DEPS, c) were determined. Averages ± SD are shown for N=5 (A.t.: all; S.l,: WT, *slkea3-2*) or N=6 (S.l.: *slkea3-1*). A Kruskal–Wallis test followed by Dunn’s post hoc test was applied for statistical analysis. Asterisks indicate significant differences to Col-0. ns, not significant.

Because KEA3 activity was shown to affect the de-epoxidation state of the xanthophyll pigments under fluctuating light conditions (von Bismarck *et al*., 2023), we analyzed the levels of xanthophyll cycle pigments (VAZ pool) and their de-epoxidation state (DEPS, Fig. 6b-c, Fig. S13) in both species. The VAZ-to-chlorophyll ratio is markedly higher in tomato than in *A. thaliana*. In line with the previous report, *atkea3* mutants displayed a significant increase in DEPS compared with Col-0, while both *slkea3* mutants showed only a slight, non-significant rise in DEPS compared with M.M. (Fig 6b, c). WT plants of the two species had comparable DEPS levels.

### Species-specific computational models recapitulate NPQ dynamics and effects of KEA3

Attempts to replace AtKEA3 in *A. thaliana* with SlKEA3 to investigate possible effects of the C-terminal extension on KEA3 regulation and thus NPQ dynamics were unsuccessful. Hence, we followed a computational approach to dissect the differences in KEA3-dependent NPQ relaxation. An ordinary differential equation (ODE) model for NPQ was established based on previous models ((Matuszyńska *et al*., 2016; Li *et al*., 2021; Seydoux *et al*., 2022); see Fig. S14 for summarizing scheme and Method S1 for a full model description). The new model included the following two main adjustments: (i) changes in the contribution of PsbS and zeaxanthin to NPQ, and (ii) the inhibition of KEA3 activity by ATP and stromal pH. The best-fitting sets of parameters for the species-specific models were identified by Monte Carlo simulations (Fig S15, Table S4). The final models recapitulated the NPQ dynamics of WT and *kea3* mutants in both species during light fluctuations (Fig. 7a, b). A limitation of the model was apparent during the first transition from darkness to low light, in which NPQ was under-estimated in *A. thaliana* (Fig. 7b, Fig. S16). However, the model performed well under recurrent fluctuations.

**Figure 7.**
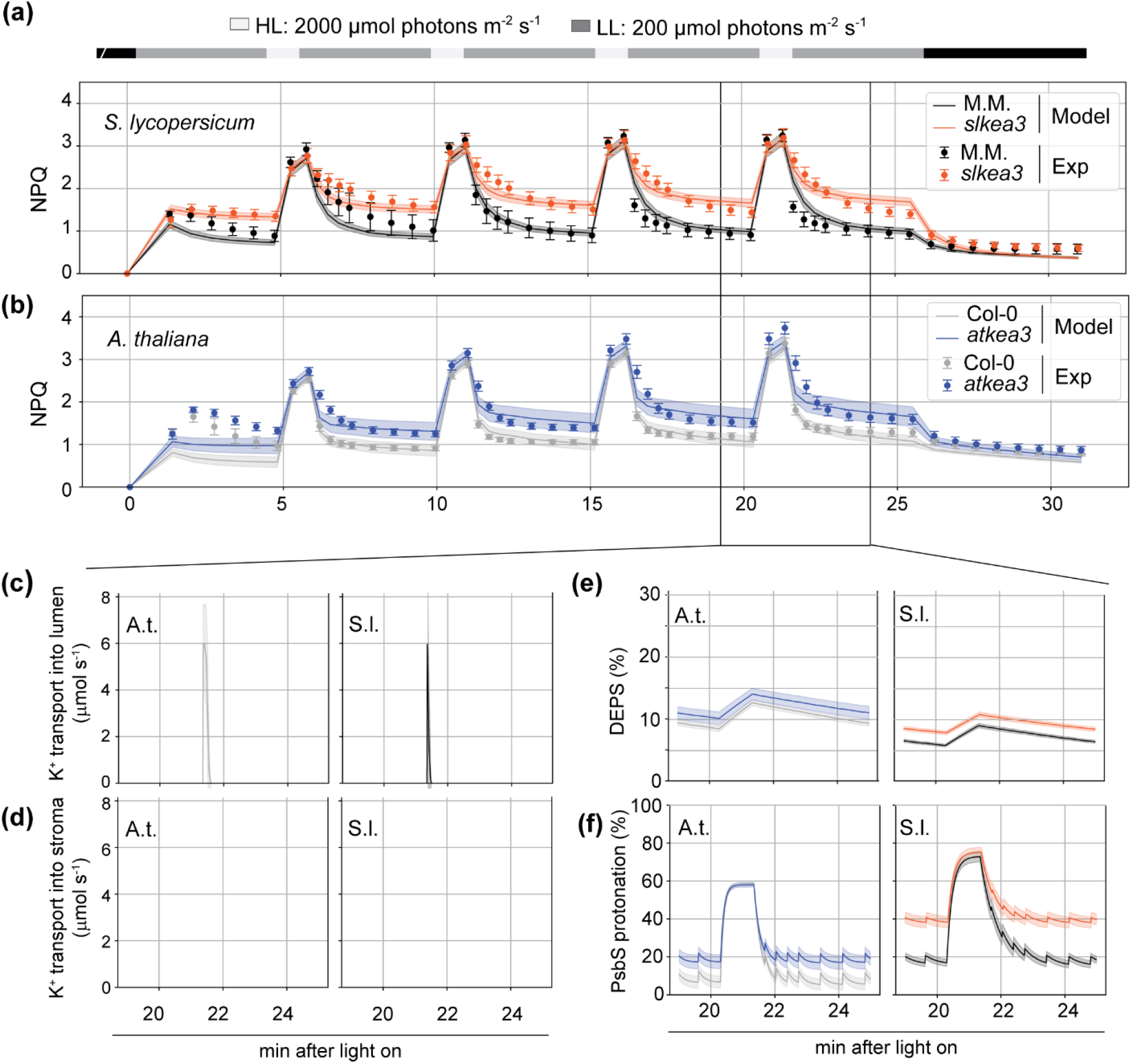
A kinetic mathematical model reproduces dynamic NPQ during light fluctuations for both species. (a-b) The model was fitted simultaneously to the average NPQ of WT and of the two *kea3* mutants from *S. lycopersicum* (a) and *A. thaliana* (b) as shown in Fig.6. (c-f) KEA3-mediated K^+^ transport from the stroma into lumen (c) and from the lumen into the stroma (d), the de-epoxidation state (DEPS) of the xanthophylls (e) and PsbS protonation (f) during the fourth light fluctuation..

The KEA3 in the model does not mediate K^+^ outflux, but only influx, and it is only transiently active after a high to low light transition (Fig. 7c-d, Fig. S16). The best-fitted parameter for ATP inactivation was higher in *A. thaliana* than in the tomato model (Table S3), which is in line with differences in regulation due to structural variations in the regulatory stromal C-terminus. KEA3 activity started 26.3 ms after the transition from high to low light in both models (4^th^ fluctuation), but the duration of activity was ∼10x longer in *A. thaliana* (135.2 ms) than in tomato (11.3 ms). The model predicted KEA3 activity does not affect the PMF size but increases the contribution of ΔΨ similarly in both species (Fig. S17). The resulting change in PMF partitioning, and thud lower ΔpH reduced NPQ.

The two main components of qE, the PsbS protein and the xanthophyll cycle, showed species-specific features from the model’s predictions (Figure 7 e-f, Table S3). DEPS was a little higher in *kea3* mutants of both species than in WT, slightly increasing and decreasing during the high and low light phase, respectively (Fig. 7e). However, the dynamics (i.e., the amplitude between low and high light) in DEPS over the flight fluctuations were small compared to the PsbS protonation state. Differences in PsbS kinetics were much larger in tomato than in *A. thaliana*, with a lack of KEA3 resulting in higher PsbS protonation during the low light phases in this species with major effects on NPQ in this model (Fig. 7f).

Summing up, the main variation in the effect of KEA3 on NPQ dynamics between tomato and *A. thaliana* was predicted to result from differences in PsbS protonation rather than from differences in KEA3 activity or regulation.

### PsbS sequences from *A. thaliana* and tomato differ in surface charges surrounding one pH sensing glutamate

The model prediction of marked differences in PsbS protonation in low light prompted us to have a closer look at the PsbS protein sequences of the two species with specific focus on the vicinity of the two protonatable and thus pH sensing glutamates. Indeed, the PsbS from the tomato and *A. thaliana* cultivar/accessions used in this study differ in the amino acid adjacent to the second pH-responsive glutamate (isoleucine in *A. thaliana* and phenylalanine in tomato; Fig. 8a). Predictions by Alphafold3 showed overall very similar protein structures for PsbS from both species, particularly of the four transmembrane helices and the lumen facing side. However, the structures differ in the orientation of the glutamate side chain and the surface charge of the encompassing region (Fig. 8b, Fig. S18). These differences may indeed affect the dynamics of the protonation of the second glutamate specifically.

**Figure 8.**
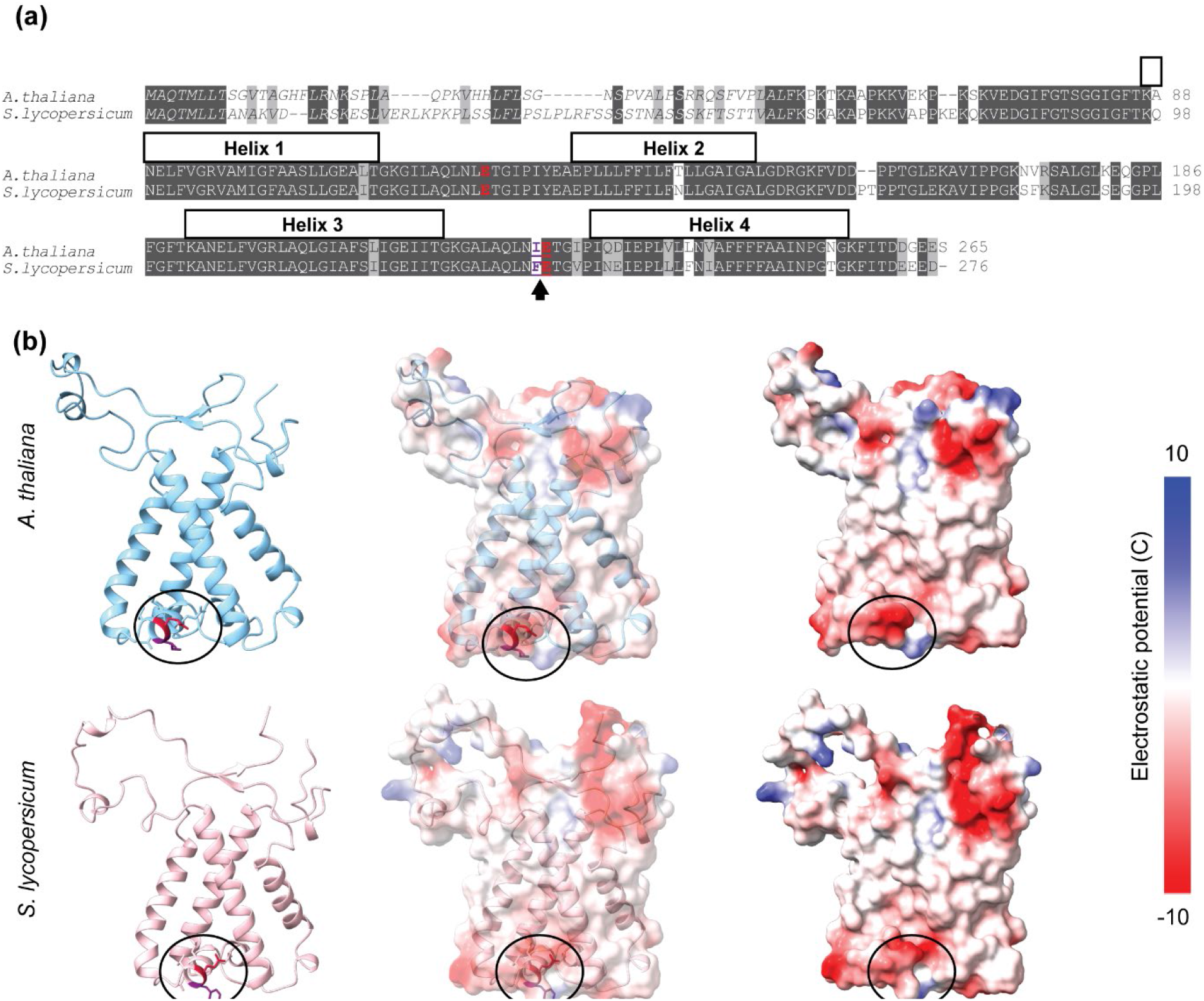
Arabidopsis and tomato PsbS differ in an amino acid adjacent to one pH-sensing glutamate with predicted effects on surrounding surface charges. (a) Alignment of PsbS protein sequences from *A. thaliana* and tomato (*S. lycopersicum*) showing the pH sensing glutamates (E) in bold and red. Identical amino acid residues are marked with a dark grey background, similar ones in light grey. The protein sequences differ in the amino acid N-terminal of the second glutamate, marked in bold and purple and highlighted by an arrow. (b) Protein structure predicted by Alphafold 3 (left) shows differences in the orientation of the glutamate side chain (red) and position of adjacent the isoleucine (A.t.) or phenylalanine (S.l) coloured in purple. The right panel shows the surface electrostatic potential according to the colour scheme shown below, while the middle panel shows an overlay of predicted ribbon structure and electrostatic surface.

## Discussion

### KEA3-dependent differences in NPQ relaxation have no effect on biomass or yield of the vegetable crop tomato

In recent years, much effort has been directed toward enhancing crop yields by modifying photosynthesis (Croce *et al*., 2024; Long *et al*., 2025). One prominent strategy has focussed on accelerating NPQ relaxation to increase light-energy conversion and, consequently, yield. The resulting growth and yield effects have varied among species, with some showing benefits and others deficits (Kromdijk *et al*., 2016; Garcia-Molina & Leister, 2020; De Souza *et al*., 2022; Lehretz *et al*., 2022; Zheng *et al*., 2026). Part of this variation may stem from the specific VPZ construct used across species; the overexpressed ZEP is also a key enzyme in the ABA biosynthesis pathway (Koornneef *et al*., 1982; Niyogi *et al*., 1998; Park *et al*., 2008). In VPZ-expressing potato lines, which exhibited reduced growth and yield under fluctuating light, elevated ABA levels were reported (Lehretz *et al*., 2022). We therefore pursued the modulation of NPQ relaxation kinetics in a crop species by manipulating KEA3 protein abundance, a strategy expected to avoid secondary effects on plant traits via phytohormone changes.

Our results show that KEA3 strongly accelerated NPQ relaxation in tomato grown under commercial (polytunnel) conditions. Yet, differences in NPQ relaxation between genotypes were only weakly linked to CO_2_ assimilation and did not translate into consistent changes in growth or yield (Fig. 2-3). The results point to multiple layers of buffering between NPQ relaxation kinetics, photosynthetic electron transport, CO_2_ assimilation, and ultimately biomass production and yield. KEA3-deficient tomato lines relaxed NPQ very slowly and maintained higher NPQ throughout low-light periods (Fig. 2, Fig. S3-5). In the PT1 trial, NPQ and Φ_PSII_ were inversely correlated across all genotypes; in the PT2 trial this relationship was evident only for the knockout mutants, suggesting that experimental differences modulated the KEA3-dependent link between NPQ and Φ_PSII_. Because the climate differed markedly between the two trials, we attributed these discrepancies to environmental factors and turned to climate-controlled RL conditions. Under these conditions, *slkea3* mutants consistently displayed elevated NPQ and a transient drop in Φ_PSII_ after high to low light transitions, but CO_2_ assimilation was only minimally reduced relative to WT (Fig. 2-3, Fig. S7). During low light phases, A_net_ dropped to ∼80% of WT for less than 40 s following a high light fleck (Fig. S7). Even such modest, short-lived differences would be expected to affect growth under strongly fluctuating light if assimilated CO_2_ were directly converted into biomass.

Together, our data suggest that tomato growth and yield is not affected by NPQ relaxation kinetics, at least not under the tested conditions. The weak link between NPQ and CO_2_ assimilation may in part be due to photosynthetic limitations, such as stomatal conductance, Rubisco activity, or triose-phosphate utilization (reviewed by (McClain & Sharkey, 2019; Croce *et al*., 2024; Long *et al*., 2025). The missing translation from CO_2_ assimilation to growth could involve source-sink interactions (reviewed by (Paul *et al*., 2020; Rosado-Souza *et al*., 2023)) or external factors like pot size (Poorter *et al*., 2012), soil properties, or water availability.

### KEA3 shows little activity under drought stress

In the PT2 trial, temperatures exceeded 40°C around the time of the measurements (Fig. S2), and the watering regime had to be adjusted to match the increased requirement. Transient water and heat stress likely explains why the effects of KEA3 levels on NPQ and Φ_PSII_ was slightly less pronounced in PT2 than PT1 (Fig. 2). Indeed, when we analyzed the NPQt/ECSt ratio as a proxy for KEA3 activity in control and drought-stressed plants growing under climate-controlled RL conditions, we found differences in *slkea3* mutants compared to WT only in the control but not in the drought-treated plants, supporting that KEA3 activity is repressed under water deficit (Fig. 4). That the reduced watering regime caused drought stress was confirmed by a markedly reduced plant size and altered ion content (Fig. 4e-f, Fig. S8-9, also reported in (D’Oria *et al*., 2022; Permatasari *et al*., 2026)). The inhibition of KEA3 under drought can be explained in line with metabolic regulation of its activity: during drought, reduced turnover of ATP and NADPH in the carbon reactions is sensed by the stromal regulatory domain which gates K^+^/H^+^ exchange (Uflewski *et al*., 2021, 2024). Because NPQ is protective under drought, inactivation of KEA3 is required. Accordingly, we did not observe any effect of a lack of KEA3 on drought resistance in tomato (Fig. 4e-f). These results contrast with recent work in *A. thaliana*, where KEA3 was reported to be required for ABA-induced ROS production, triggering stomatal closure in drought stress, with *atkea3* mutants being more drought-susceptible than WT (Yamada *et al*., 2024).

### Combined experimental and computational analysis reveals the interaction of multiple components in species-specific photosynthetic responses

The absence of KEA3 slowed NPQ relaxation more strongly in tomato than in *A. thaliana*, yet it had a weaker impact on xanthophyll de-epoxidation (Fig. 6). Despite the higher DEPS in *atkea3*, *slkea3* exhibited a substantially stronger effect on NPQ relaxation relative to its WT. The VAZ pool size, which also influences NPQ dynamics (Johnson *et al*., 2008), was unchanged by KEA3 loss in either species, together indicating that neither differences in VAZ nor DEPS can explain the stronger effect of KEA3 on NPQ relaxation in tomato. To explore the species-specific differences systematically, we constructed a computational model replicating NPQ and Φ_PSII_ of WT and *kea3* mutants under fluctuating light. The model showed that KEA3 fluxes are much smaller than those of the photosynthetic light reactions and ATP synthase, which dominate proton exchange across the thylakoid membrane (Fig. S17).

Predicted differences in the duration of KEA3 activity are consistent with sequence and structural differences in the regulatory C-terminus (Fig. 5). However, the largest modelled effect arose from KEA3-mediated changes in PsbS protonation, which did not originate from differences in lumen pH between species (Fig. 7, Fig. S17): the model predicts that, during a low-light period, more PsbS is protonated and thus active in *slkea3* than in *atkea3* (which both have a slightly lower lumen pH than their WTs). This can be explained by a difference in the pH-dependent PsbS activation threshold between both species. Recent work demonstrated that subtle variations on the lumen-pH-sensing side of PsbS strongly affect NPQ kinetics (Chen *et al*., 2025; Mao *et al*., 2026). Notably, PsbS sequences from *A. thaliana* and tomato differ in the amino acid residue directly adjacent to the second pH-sensing glutamate (Fig. 8a). Structural and surface-charge modelling indicate that this substitution changes glutamate side chain orientation and local charge (Fig. 8b). Such differences in surface charges can affect the protonation/ deprotonation rate and tune pH sensitivity of proteins (Zhou & Pang, 2018).

## Conclusion

Our results show that differences in NPQ relaxation in *S. lycopersicum* are largely buffered across the cascade of molecular and functional steps that link this process to biomass accumulation. Although SlKEA3 strongly accelerates NPQ relaxation, its effect is progressively attenuated: first at the level of PSII quantum yield, then in CO_2_ assimilation, and finally through a large “black-box” at the level of growth and yield, where consistent measurable differences between WT and mutants were lacking. Thus, we show that a pronounced NPQ-relaxation phenotype does not necessarily translate into changes in biomass or yield, even when plants are grown under fluctuating or natural dynamic light environments. By integrating the experimental data from tomato and *A. thaliana*, we constructed a computational model that accurately recapitulates species-specific NPQ dynamics and uncovers molecular variations underlying these distinct responses. The model therefore provides mechanistic insights that could not be obtained from the individual phenotypes alone. While it does not link NPQ kinetics directly to field productivity, it offers a framework that can be leveraged for future crop-enhancement strategies, once the dominant field-level constraints that limit translation from NPQ dynamics to yield are identified.

## Data availability

The model was implemented in Python using the mxlpy library (van Aalst *et al*., 2025). Model code, parameter sets, and simulation scripts are available as a public repository on the ARC Datahub (https://git.nfdi4plants.org/zat55vor/kea3_tomato). An interactive implementation allowing direct exploration of model dynamics is available through mxl-web (https://computational-biology-aachen.github.io/mxl-web/models/kea3-tomato). All raw data and analyses can be found on ARC Datahub (https://git.nfdi4plants.org/zat55vor/kea3_tomato).

## Author contributions

UA and TVB designed the plant experiments. TVB performed all plant experiments together with JR, LS and JD, except for the pigment analysis done by QHN and PJ and the ionome analysis done by MK and HHK. KK helped with experimental design and monitored environmental parameters. QHN and AM constructed the mathematical model. UA analyzed protein structure predictions. TVB, QHN and UA wrote the manuscript with input from all authors.

## Acknowledgement

We thank K. Lepa-Klabe for performing the tomato transformations, D. D. Zerning and L. Westphal for plant maintenance in the polytunnel experiments, S. Bethmann for running the HPLC, D. C. Velazquez Ayala for help with MultispeQ measurements; D. Vrbic for setting up the openJII workbook; J. Ebeling for critical input and comments on the model. The work was funded by 02Flux4LIVES (DFG 355293141, H-H. K., U. A.), CEPLAS (EXC 2048/1–390686111, Q.H.N, A.M., U. A) and FOR 5573 (DFG 507704013, H-H. K., A.M, U.A.)

