## Supplementary material for "Experimental and computational analyses of KEA3-dependent photosynthetic responses uncover species-specific molecular variation": Fig. S

The following Supporting Information is available for this article:

**Fig. S1** **Immunoblot analysis probing KEA3 content and HA-tag of *oeSlKEA3*.**

**Fig. S2** **Climate data recorded during polytunnel experiments.**

**Fig. S3** **The effect of KEA3 abundance and growth condition on NPQ and $\Phi_{PSII}$ dynamics.**

**Fig. S4** **Leaf age- and attachment status-dependency of NPQ and $\Phi_{PSII}$ during light intensity fluctuations of PT2-grown plants.**

**Fig. S5** **Heatmaps of KEA3 effects on NPQ and $\Phi_{PSII}$ in leaves of different developmental stage and attachment state.**

**Fig. S6** **The influence of KEA3 level on $CO_2$ assimilation during light intensity fluctuations.**

**Fig. S7** **The influence of KEA3 level on $CO_2$ assimilation during light intensity fluctuations. Changes in NPQ and $CO_2$ assimilation rate in RL.**

**Fig. S8** **Analysis of the ionome of control and drought treated plants does not reveal any differences between M.M. and the knock-out lines.**

**Fig. S9** **Analysis of the transition metal ionome of control and drought treated plants does not reveal any differences between M.M. and the knock-out lines.**

**Fig. S10** **The C-terminal extension forming a β-sandwich is present in all analyzed *Solanum* species.**

**Fig. S11** **Alphafold-models of KEA3 dimers with confidence scores.**

**Fig. S12** **Comparison of NPQ and $\Phi_{PSII}$ responses under light fluctuations between *A. thaliana* and tomato.**

**Fig. S13** **VAZ pool pigments from *A. thaliana* and tomato.**

**Fig. S14** **Scheme illustrating the construction and components of the mathematical model.**

**Fig. S15** **Monte Carlo simulation on chosen parameter sets.**

**Fig. S16** **Modelled dynamics of selected photosynthetic parameters during light intensity fluctuations.**

**Fig. S17** **Modelled dynamics of selected KEA3-related parameters during light intensity fluctuations.**

**Fig. S18** **PsbS structure predictions and surface electrostatic potential.**

**Methods S1 Mathematical model for NPQ kinetics in *tomato* and *A. thaliana***

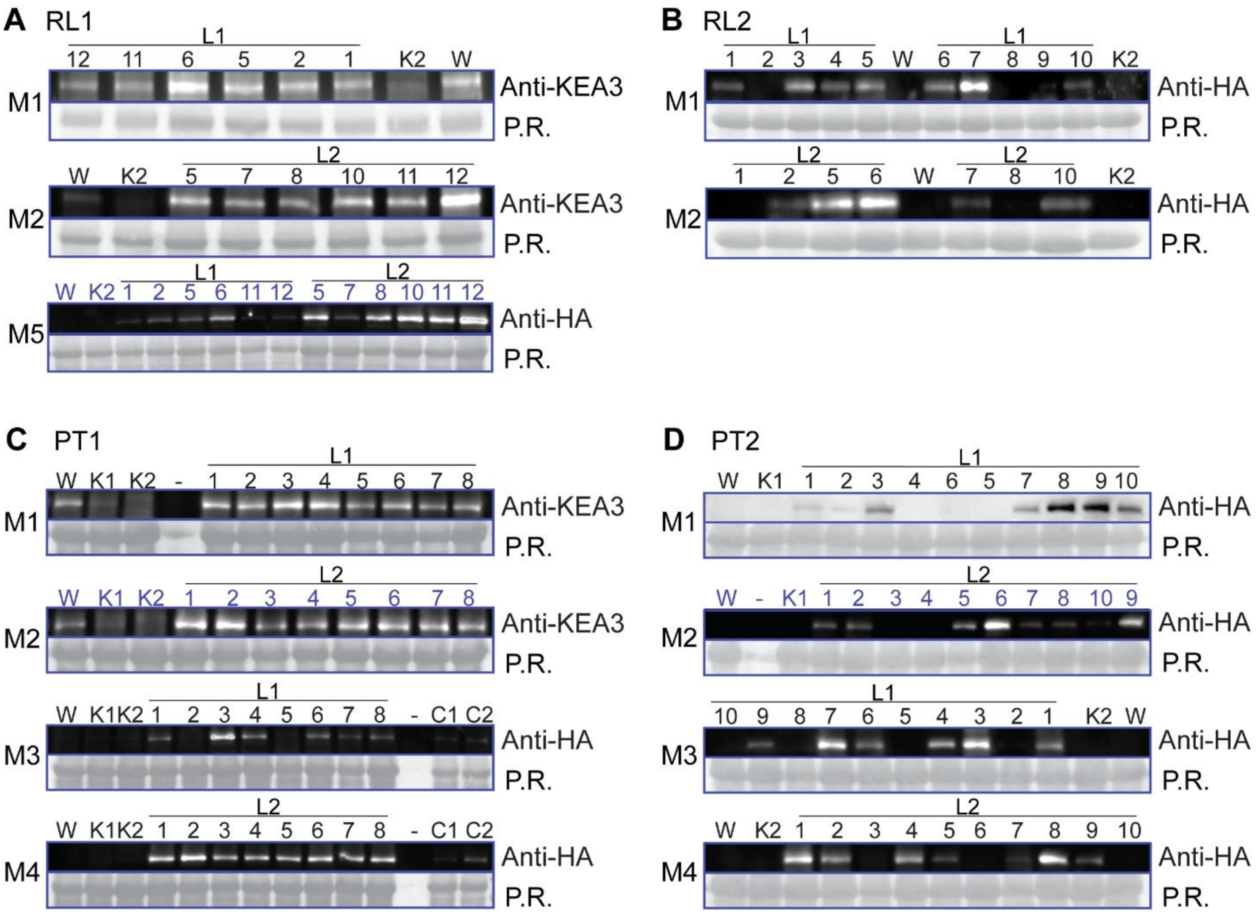


**Fig. S1 Immunoblot analysis probing KEA3 content and HA-tag of *oeSlKEA3*.** Total leaf extracts were used and membranes were incubated either with an N-terminal KEA3-antibody (anti-KEA3) or an HA-antibody (anti-HA). The 55 kDa band stained with ponceau red (P.R.) corresponding to the large subunit of rubisco is shown as loading control. Numbers (replicate of *oeSlKEA3-L1* or *L2*) and letters (W – Wildtype, K1 – *slkea3-1*, K2 – *slkea3-2*, C1 – Positive control 1 for HA, C2 – Positive control 2 for HA) above each lane indicate the loaded genotype and replicate number. For PT2, two separate sets of plants, grown in parallel, were used for all physiological measurements and fruit yield (M1-M2) and for dry weight (M3-M4). The only exception were Chl a fluorescence measurements on attached leaves that were conducted directly before determination of dry weight. Some membrane images were vertically compressed for presentation purposes. No other image manipulations were performed and raw images from uncut membranes can be found on on the ARC Datahub (https://git.nfdi4plants.org/zat55vor/kea3_tomato). M – Membrane

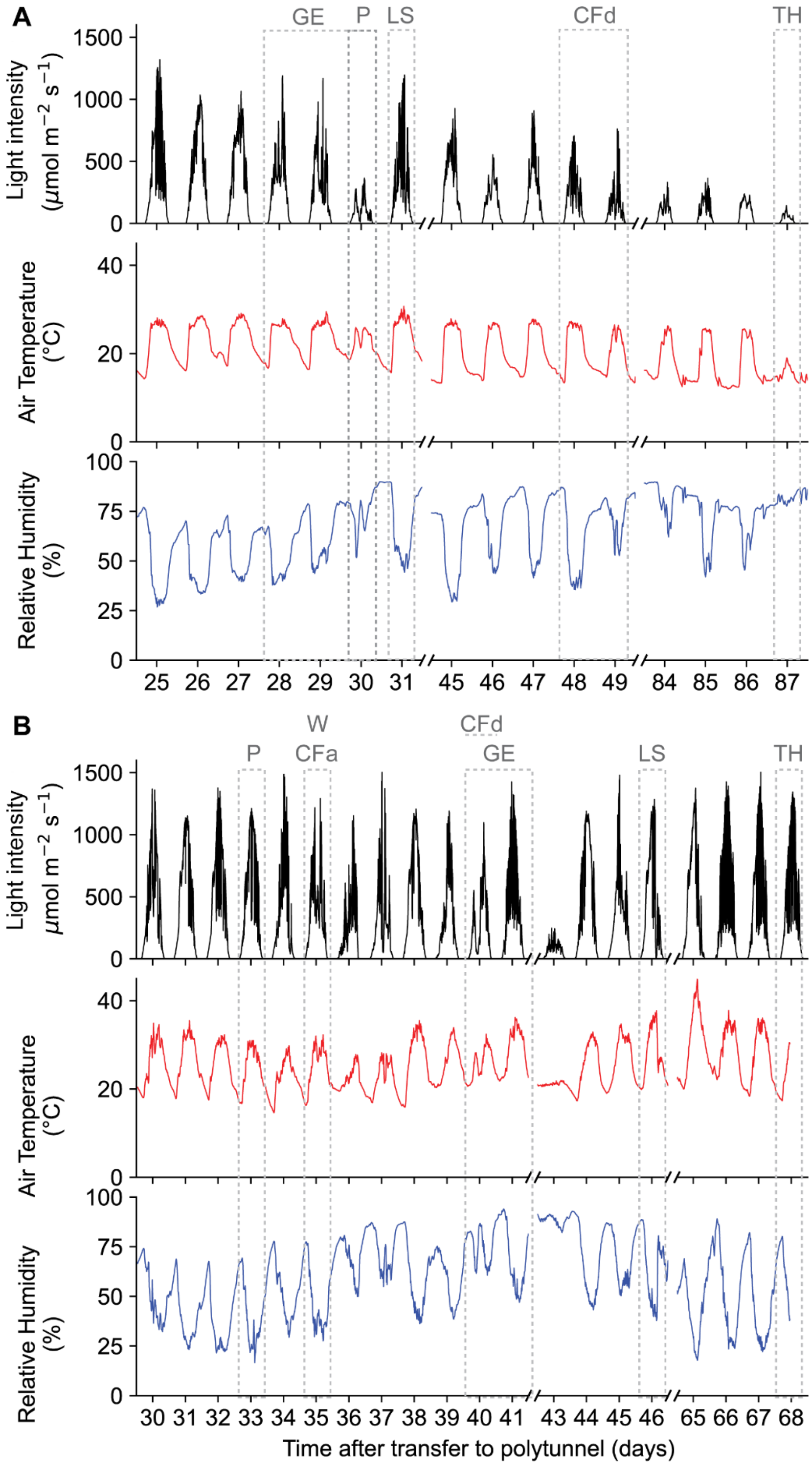


**Fig. S2 Climate data recorded during polytunnel experiments.** Light intensity (black), air temperature (red) and relative humidity is shown for the period in which measurements were performed during polytunnel experiment 1 (PT1; A) and 2 (PT2; B). Measurement days are indicated by dotted lines. Complete climate data can be found in Table S3. GE – Gas exchange measurements, P – Photos, LS – Leaf sample taken, CFd – Chlorophyll fluorescence measurements on detached leaflets, CFa – Chlorophyll fluorescence measurements on attached leaflets, TH – Tomato harvest, W – Weight measurement.

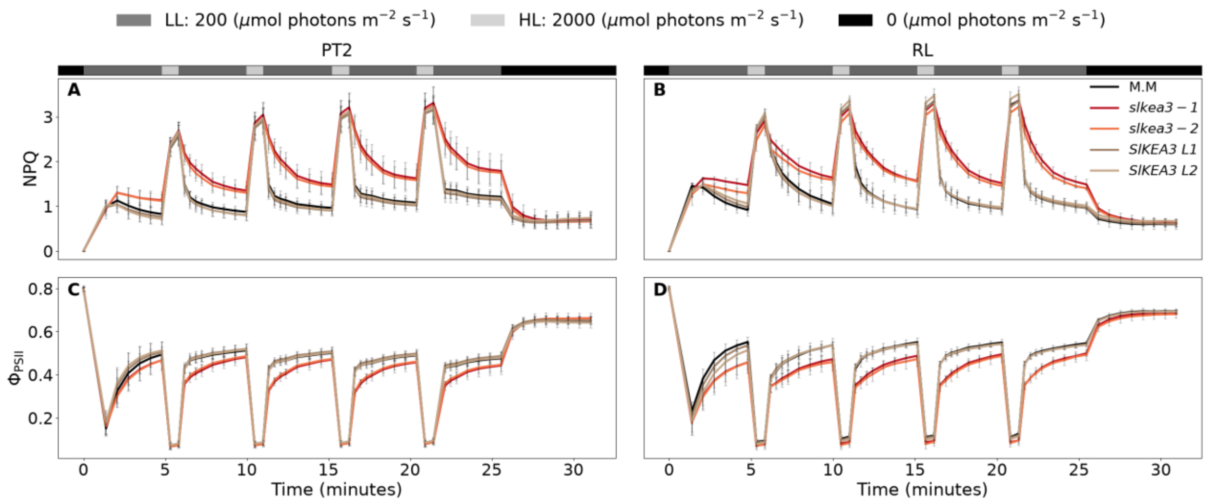


**Fig. S3 The effect of KEA3 abundance and growth condition on NPQ and $\Phi_{PSII}$ dynamics.** Chlorophyll *a* fluorescence was measured on the 30 min dark adapted 2nd or 3rd leaflet during light intensity fluctuations of 4 min low light (200 µmol photons $m^{-2}$ $s^{-1}$) and 1 min high light (2000 µmol photons $m^{-2}$ $s^{-1}$), a final 4 min low light phase and subsequent 5 min darkness. Averages ± SD of NPQ (A) and $\Phi_{PSII}$ (B) are shown with N=4 (RL: *slkea3-1*), N=5 (PT2: *SlKEA3 L1*, *SlKEA3 L2*; RL: WT, slkea3-1, *SLKEA3 L2*), N=6 (RL: *SlKEA3 L1*), N=7 (PT1: WT, *slkea3-1*, *slkea3-2*, *SlKEA3 L2*, PT2: WT, *slkea3-1*, *slkea3-2*). Selected time points were used to generate heatmaps in Fig. 2.

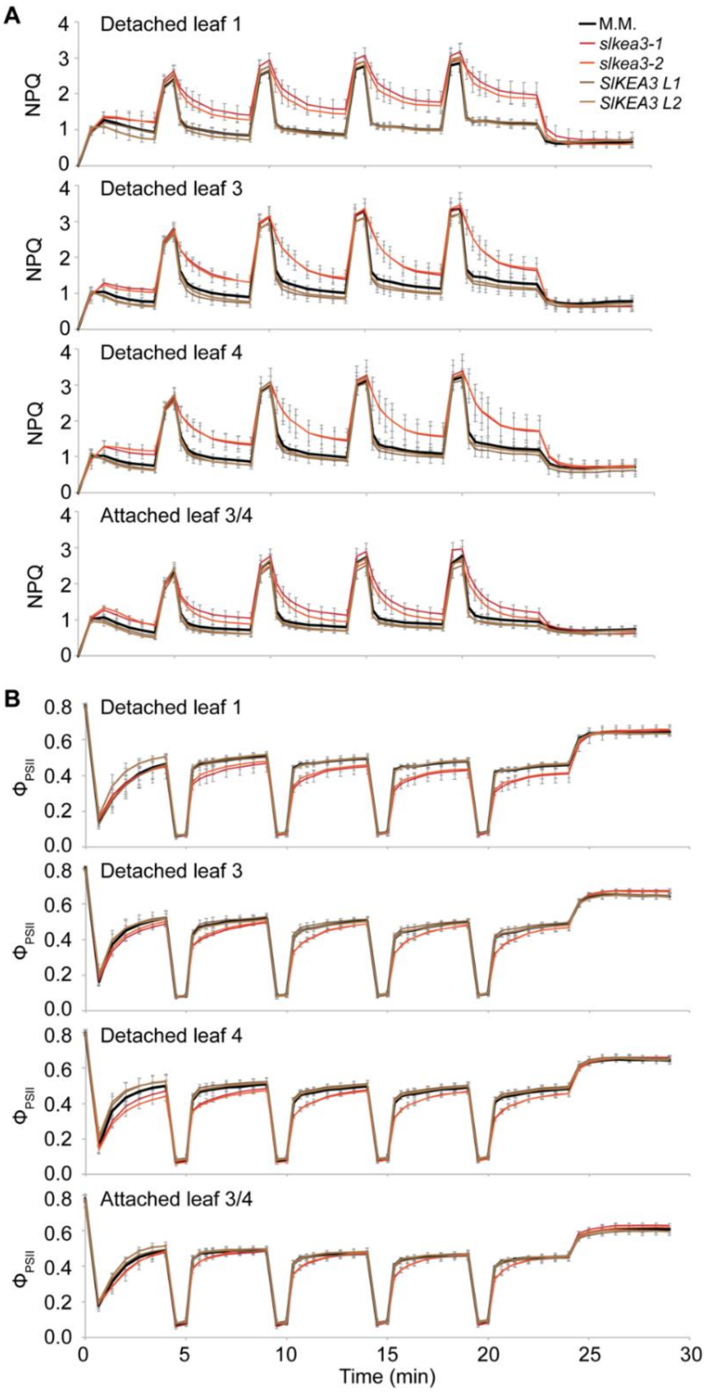


**Fig. S4 Leaf age- and attachment status-dependency of NPQ and $\Phi_{PSII}$ during light intensity fluctuations of PT2-grown plants.** Chlorophyll *a* fluorescence was measured on 30 min dark adapted, detached 2nd or 3rd leaflets of leaves of different developmental stage (counting from leaf 1 being the youngest leaf >5cm of length) and on the attached 2nd or 3rd leaflets of leaf 3 or 4 during four intensity fluctuations of 4 min low light (200 µmol photons $m^{-2}$ $s^{-1}$) and 1 min high light (2000 µmol photons $m^{-2}$ $s^{-1}$), a final 4 min low light phase and 5 min darkness. For the attached leaflet, either leaf 3 or 4 was chosen for comparable size between replicates and genotypes. Averages ± SD are shown of detached leaves with N=5 (*SlKEA3 L1*, *SlKEA3 L2*) or N=7 (M.M., *slkea3-1*, *slkea3-2*) and attached leaves with N=2 (*slkea3-2*), N=3 (*SlKEA3 L1*, *SlKEA3 L2*), N=4 (M.M) and N=5 (*slkea3-1*). Leaf 1 was defined as the youngest leaf with a length ≥ 5cm.

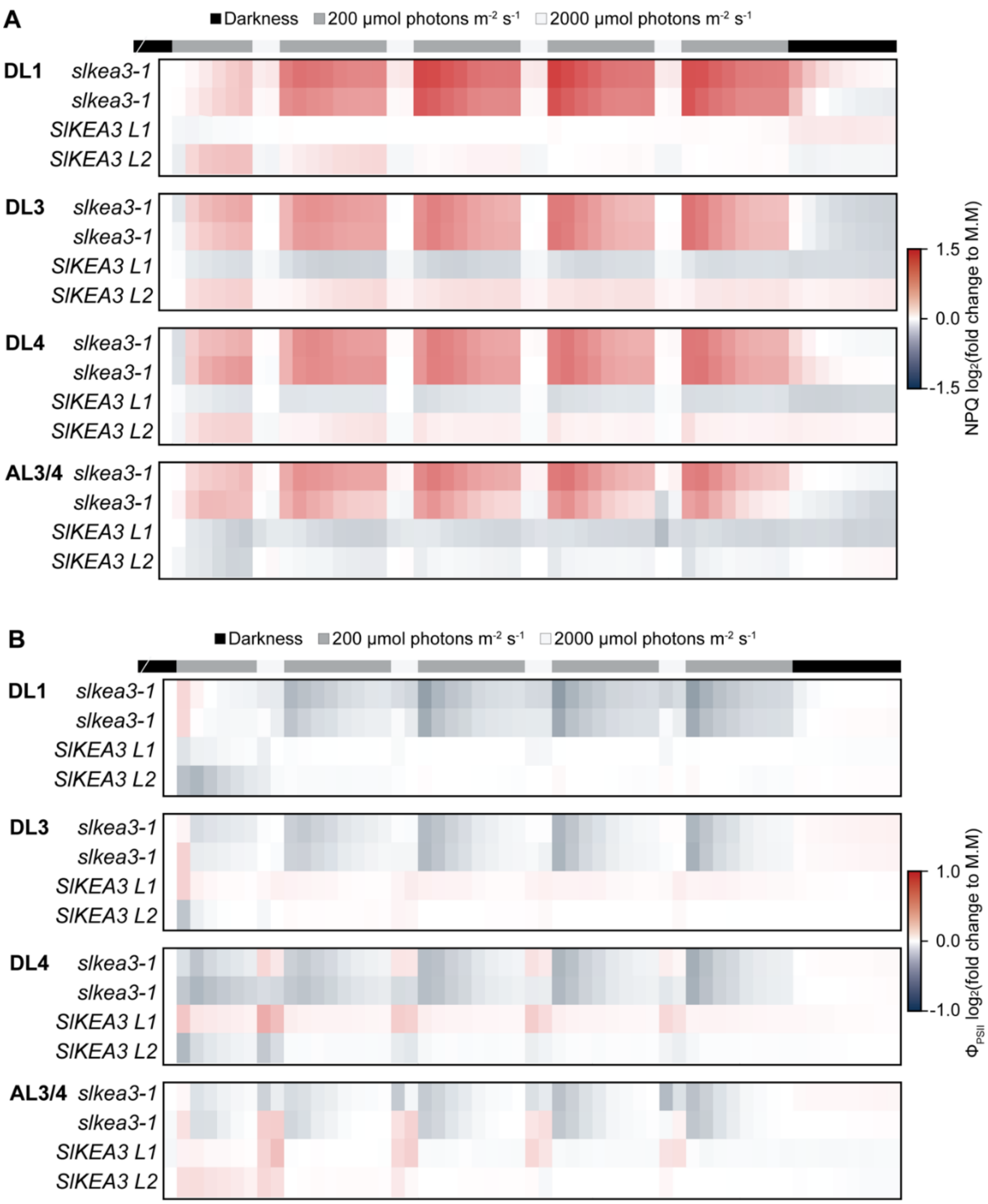


**Fig. S5 Heatmaps of KEA3 effects on NPQ and $\Phi_{PSII}$ in leaves of different developmental stage and attachment state.** Data from Fig. S2 (NPQ, A and $\Phi_{PSII}$, B) were transformed into heatmaps. The false color scale indicates log-fold changes compared to the wildtype M.M..

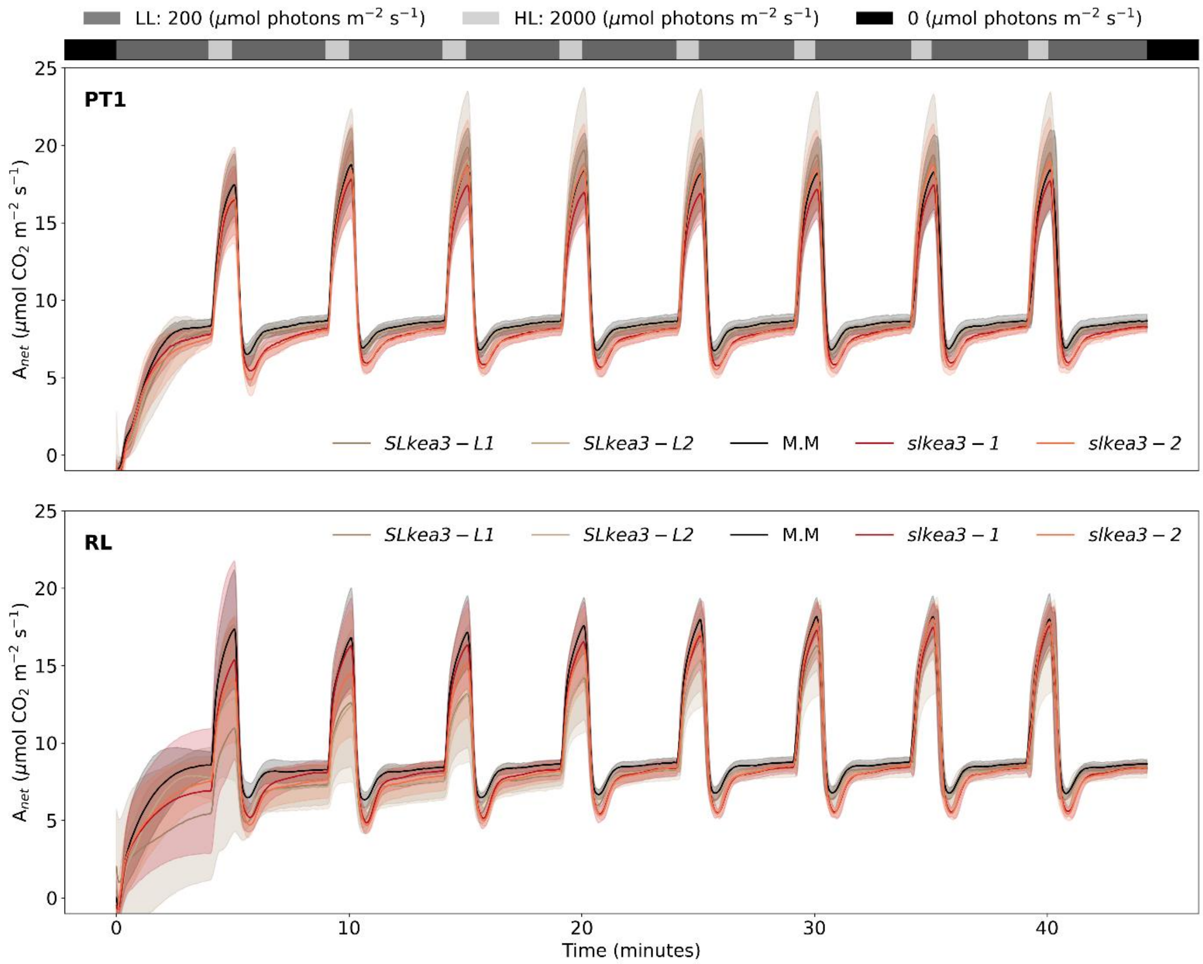


**Fig. S6 The influence of KEA3 level on $CO_2$ assimilation during light intensity fluctuations.** $CO_2$ assimilation rate ($A_{net}$) was measured on the 2nd or 3rd leaflet of leaf 4 (counting from leaf 1 being the youngest leaf >5cm of length) during eight light intensity fluctuations of 4 min low light (200 µmol photons $m^{-2}$ $s^{-1}$) and 1 min high light (2000 µmol photons $m^{-2}$ $s^{-1}$) and a final 4 min low light phase. Shown are the averages ± SD of plants grown in the polytunnel experiment 1 (A; PT1) or controlled, random fluctuating light conditions (B; RL) with N=5 (PT1: *SlKEA3 L1*, RL: SlKEA3 L1, *SlKEA3 L2*), N=6 (RL: *slkea3-1*, *slkea3-2*) and N= 8 (PT1: WT, *slkea3-1*, *slkea3-2*, *SlKEA3 L2*; RL: WT). Bins of selected time points are shown as heatmaps in Fig. 3.

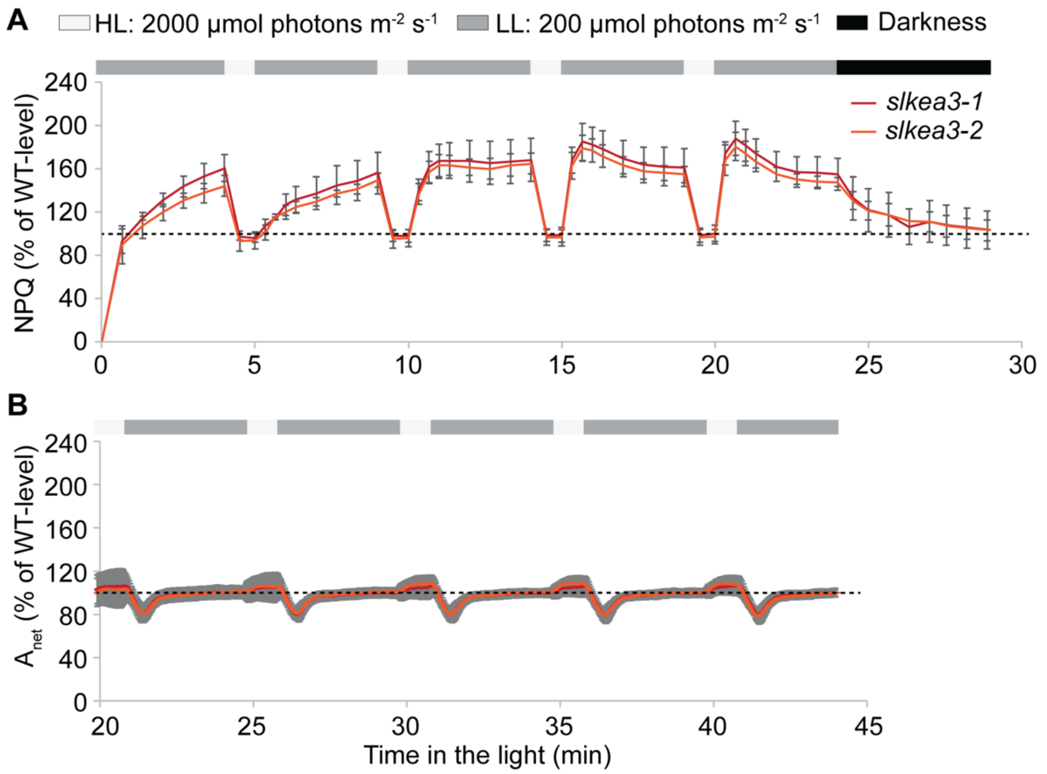


**Fig. S7 The influence of KEA3 level on $CO_2$ assimilation during light intensity fluctuations. Changes in NPQ and $CO_2$ assimilation rate in RL.** From data shown in Supplemental Fig. 2 and 6, the relative difference of RL-grown *slkea3-1* and *slkea3-2* to wildtype M.M. in the NPQ (A) and $CO_2$ assimilation rate ($A_{net}$; B) response to four or eight light fluctuations of 4 min low light (200 µmol photons $m^{-2}$ $s^{-1}$) and 1 min high light (2000 µmol photons $m^{-2}$ $s^{-1}$), a final 4 min low light phase and for NPQ additionally 5 min darkness was calculated. The full trace for NPQ and the last 25 min for $A_{net}$ are shown as means ±SD. Dotted lines indicate WT-level (100%).

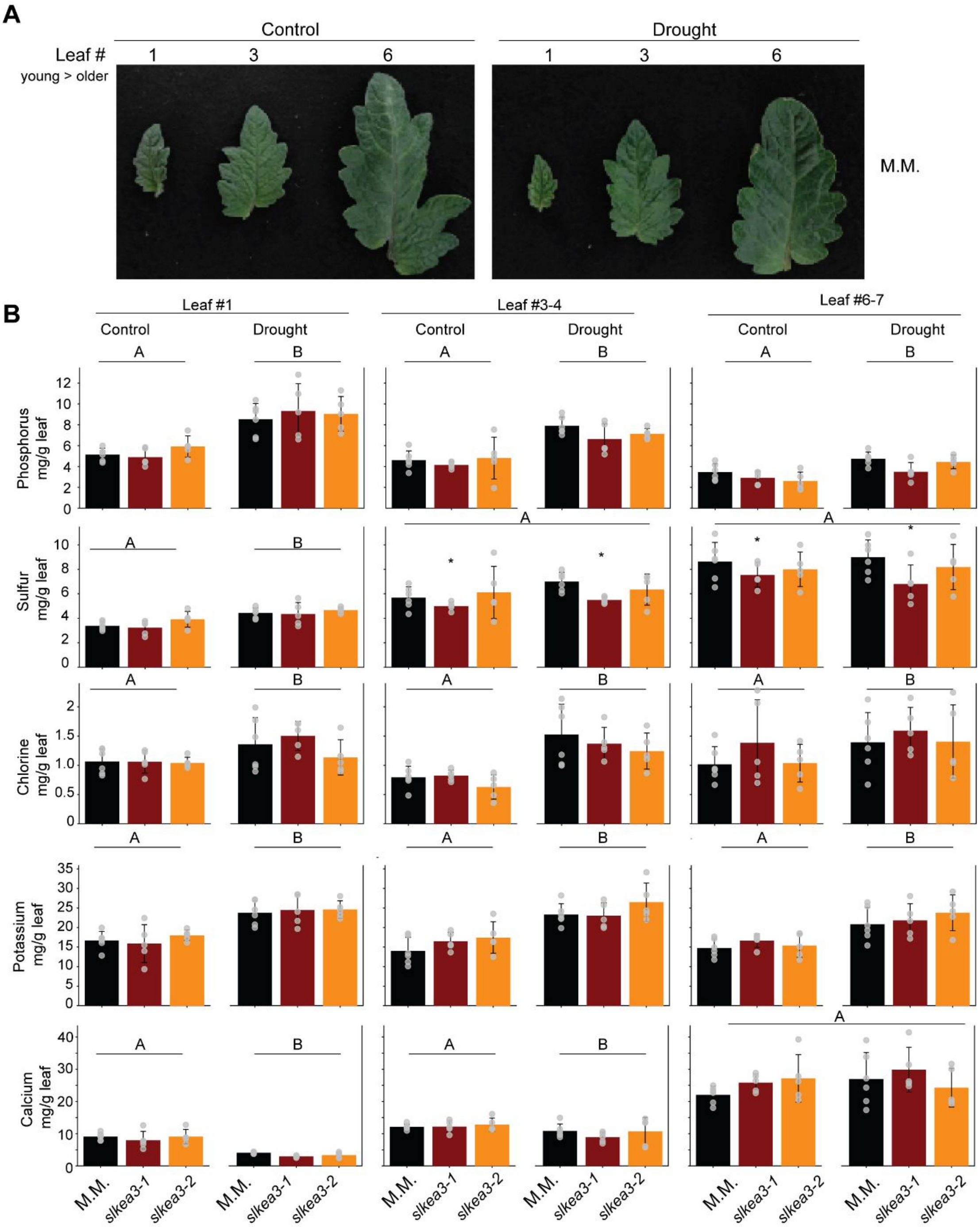


**Fig. S8 Analysis of the ionome of control and drought treated plants does not reveal any differences between M.M. and the knock-out lines.** (A) Example leaves of wildtype M.M. of different developmental stages used for analysis. Leaf#1 corresponds to the youngest leaf larger than 5 cm. (B) Drought strongly affects the levels of nutrients particularly in the younger leaves. Shown is the mean of N=6 ± SD. The statistical analysis was done by Two-way-ANOVA and Holm-Sidak multiple comparison. Capital letters A and B indicate that irrigation treatment (control vs. drought) had a significant impact on the nutrient level in the leaf, the asterisk that *slkea3-1* showed significant differences to M.M. independent of treatment.

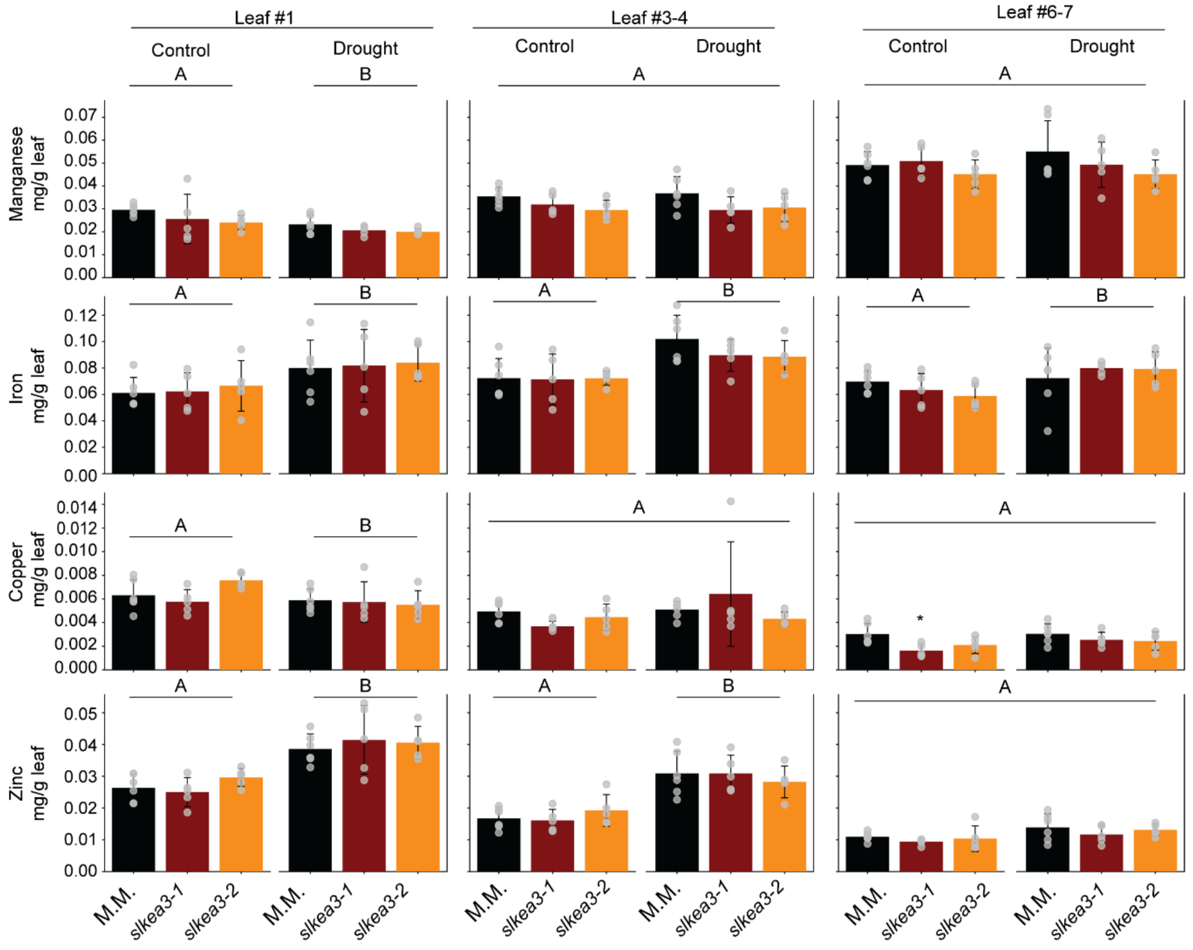


**Fig. S9 Analysis of the transition metal ionome of control and drought treated plants does not reveal any differences between M.M. and the knock-out lines**. Drought strongly affects the levels of transition metals particularly in the younger leaves. Shown is the mean of N=6 ± SD for leaf #1 (youngest leaf), #3-4 and #6-7. The statistical analysis was done by Two-way-ANOVA and Holm-Sidak multiple comparison. Capital letters A and B indicate that irrigation treatment (control vs. drought) had a significant impact on the nutrient level in the leaf.

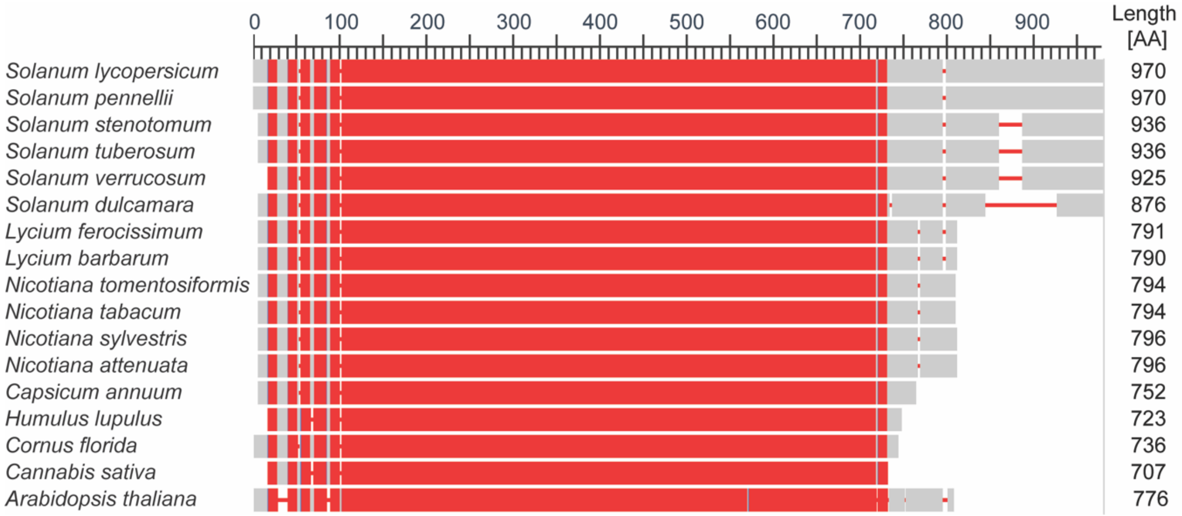


**Fig. S10 The C-terminal extension forming a β-sandwich is present in all analyzed *Solanum* species.** First 15 hits of NCBI Blast sequence comparison with the SlKEA3 amino acid sequence (plus *Arabidopsis thaliana* KEA3 sequence as comparison) show highest similarity between species of the *Solanaceae* family (first 5 hits), which all contain the C-terminal extension as compared to *A. thaliana*. Sequences were aligned using the NCBI Multiple Alignment tool, with conserved areas across all species (Conservation setting: 2 bits) colored in red.

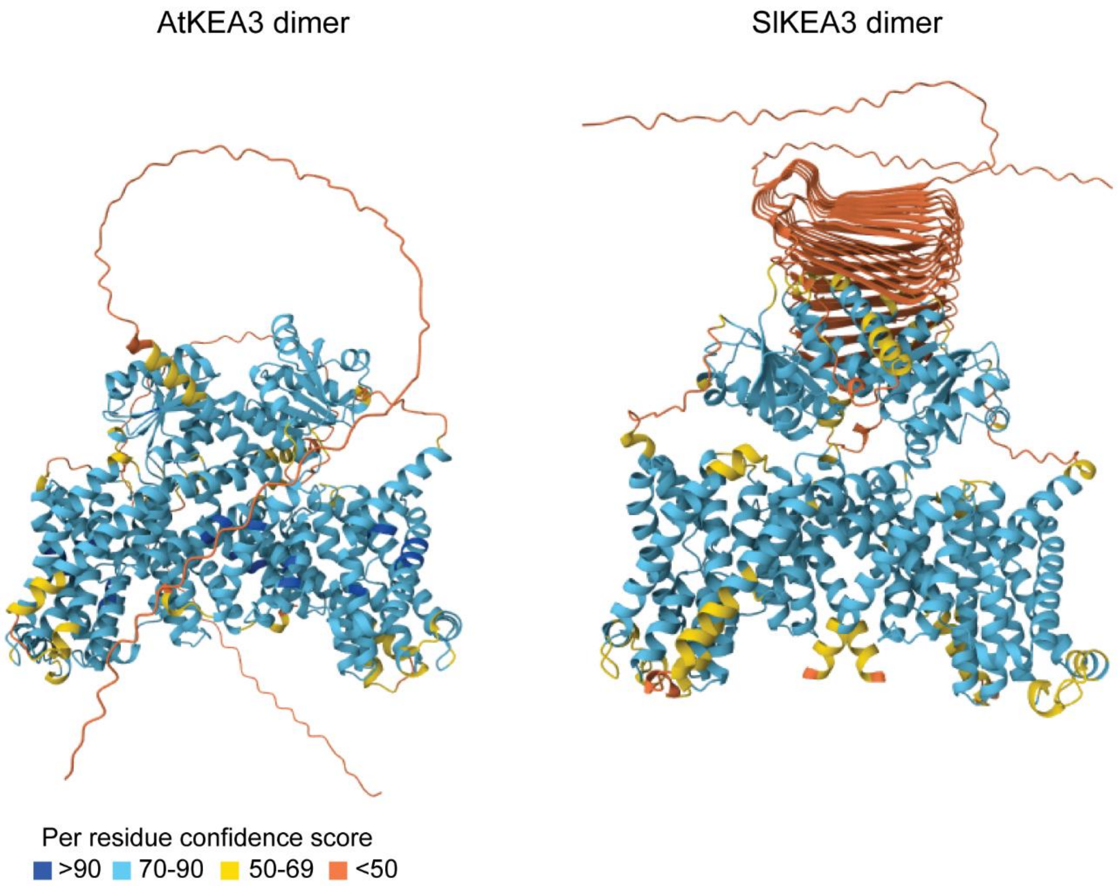


**Fig. S11 Alphafold-models of KEA3 dimers with confidence scores.** Alphafold 3 predicts homodimerization of KEA3 of both species, *A. thaliana* (left) and tomato (right). The same predictions are shown as in Fig. 5C, but with colour coding corresponding to the Alphafold confidence score of prediction.

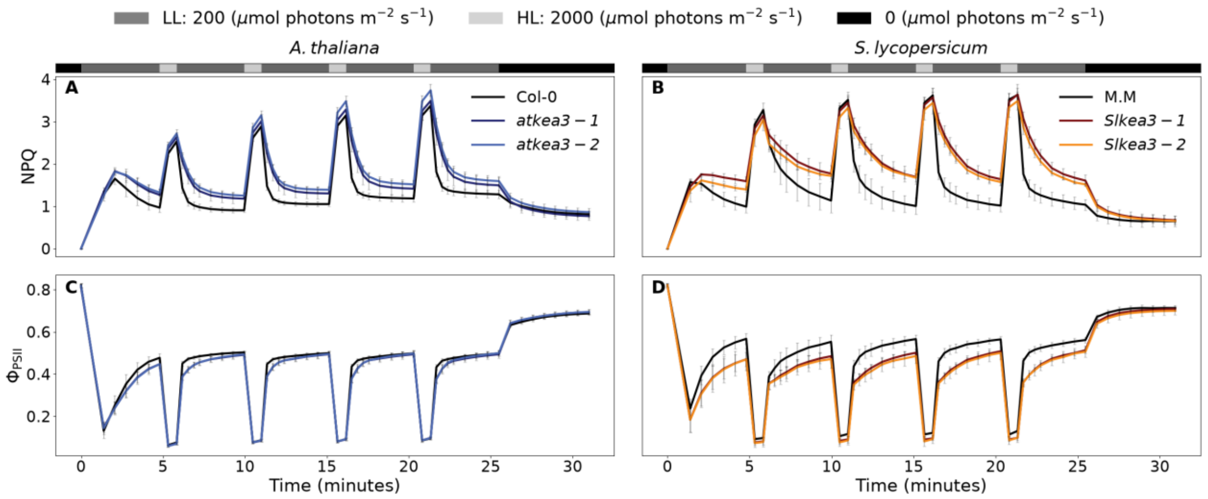


**Fig. S12 Comparison of NPQ and $\Phi_{PSII}$ responses under light fluctuations between *A. thaliana* and tomato.** Chlorophyll *a* fluorescence was measured on the 30 min dark adapted whole rosette of *A. thaliana* and 2nd or 3rd attached leaflet of leaf 3 or 4 of tomato (*S. lycopersicum*) during four light intensity fluctuations of 4 min low light (200 µmol photons $m^{-2}$ $s^{-1}$) and 1 min high light (2000 µmol photons $m^{-2}$ $s^{-1}$), a final 4 min low light phase and 5 min darkness. Averages of NPQ (A) and $\Phi_{PSII}$ (B) are shown with N=4 (S.l.: *slkea3-1*, *slkea3-2*) and N=6 (A.t.: all; S.l: M.M.) ± SD. The same data as in Supplemental Fig. 2 is shown for tomato. The NPQ response of both species during the fourth fluctuation is shown in Fig. 6.

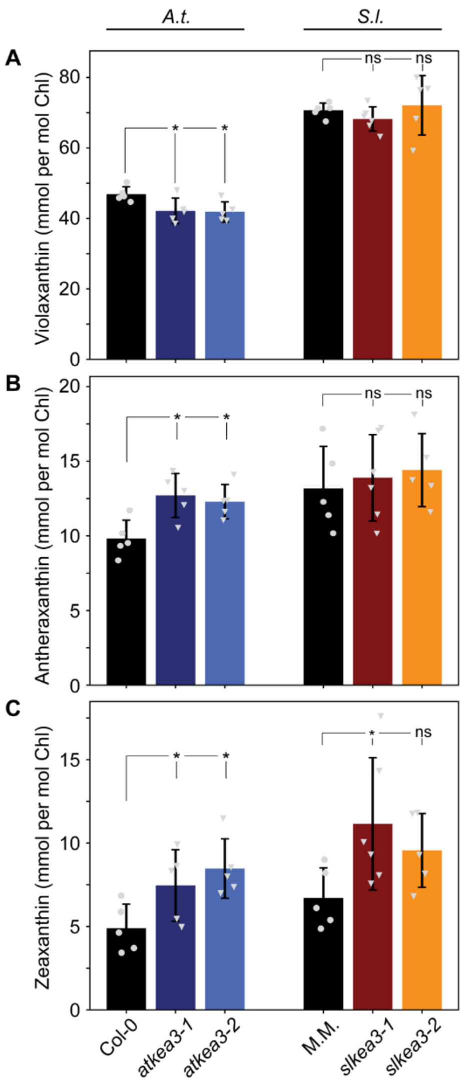


**Fig. S13 VAZ-pool pigments from *A. thaliana* and tomato.** Levels of the xanthophyll cycle pigments violaxanthin (A), antheraxanthin (B) and zeaxanthin (C) were determined via HPLC on the whole rosette (*A. thaliana*, Col-0, *atkea3-1*, *atkea3-2*) or leaves (tomato, M.M., *slkea3-1*, *slkea3-2*) collected in the middle of the light period. Shown are averages of WT and *kea3* mutants with N=5 (*A. thaliana*: all; tomato: M.M., *slkea3-2*) or N=6 (tomato.: *slkea3-1*) ± SD.

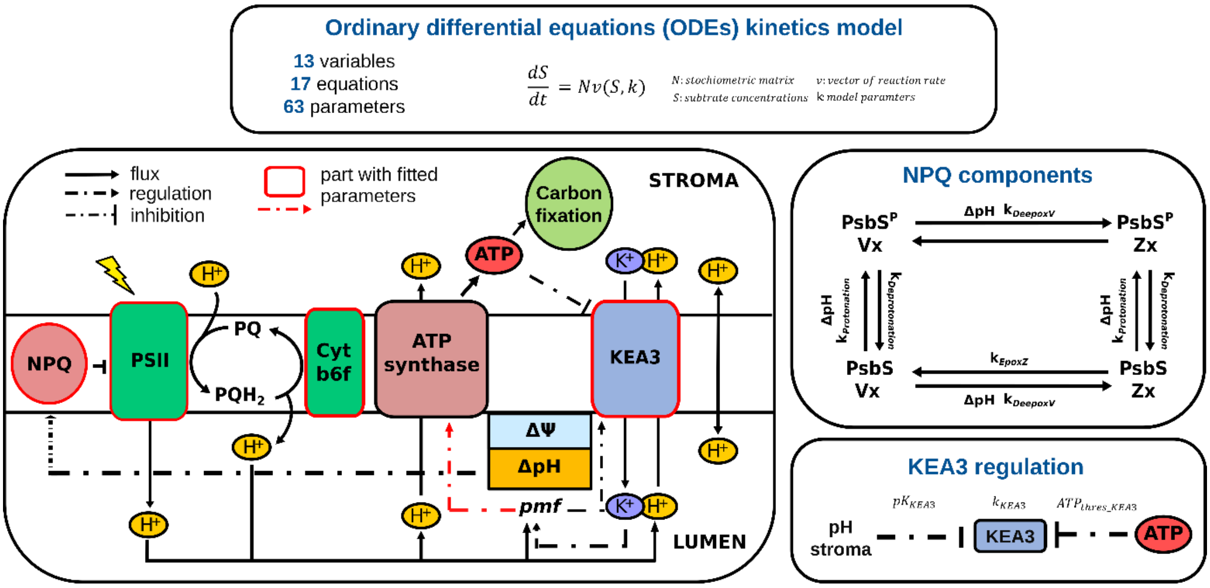


**Fig. S14 Scheme illustrating the construction and components of the mathematical model.** The model includes a simplified photosynthesis electron transport chain (PETC), the non-photochemical quenching (NPQ) process, ATP synthase activity, the proton motive force (PMF) and the $K^+$ efflux antiporter (KEA3). PETC contains photosystem II (PSII) and cytochrome $b_6f$ (Cyt b6f) activity; it is driven by light energy and translocates protons ($H^+$) into the lumen. $H^+$ accumulation in the lumen increases PMF, by contributing to the pH difference (ΔpH) and the membrane potential (ΔΨ) across the thylakoid membrane. PMF drives ATP synthesis from ADP via the ATP synthase linked to the export of $H^+$ out of the lumen. The back reaction is introduced as a simple proxy for requirements of the carbon reactions. KEA3 regulates PMF and lumen pH via coupling the efflux of $H^+$ out of the lumen with the transport of $K^+$ into the lumen. The fast NPQ components PsbS and the xanthophyll cycle pigments are activated by the low lumen pH and the subsequent increase in ΔpH. and the subsequent increase in ΔpH. PsbS can get protonated to $PsbS^P$ , while violaxanthin (Vx) will get converted into zeaxanthin (Zx). KEA3 activity is regulated via the stromal pH and ATP level, reflected through the pKa ($pK_{KEA3}$) and the ATP threshold for activation ($ATP_{thres\_KEA3}$). Within the scheme the rate constants for NPQ/KEA3-realted processes are also mentioned: PsbS protonation ($k_{Protonation}$) and deprotonation ($k_{Deprotonation}$), Vx deepoxidation ($k_{DeepoxV}$) and Zx epoxidation ($k_{EpoxZ}$), and KEA3 activity ($k_{KEA3}$).

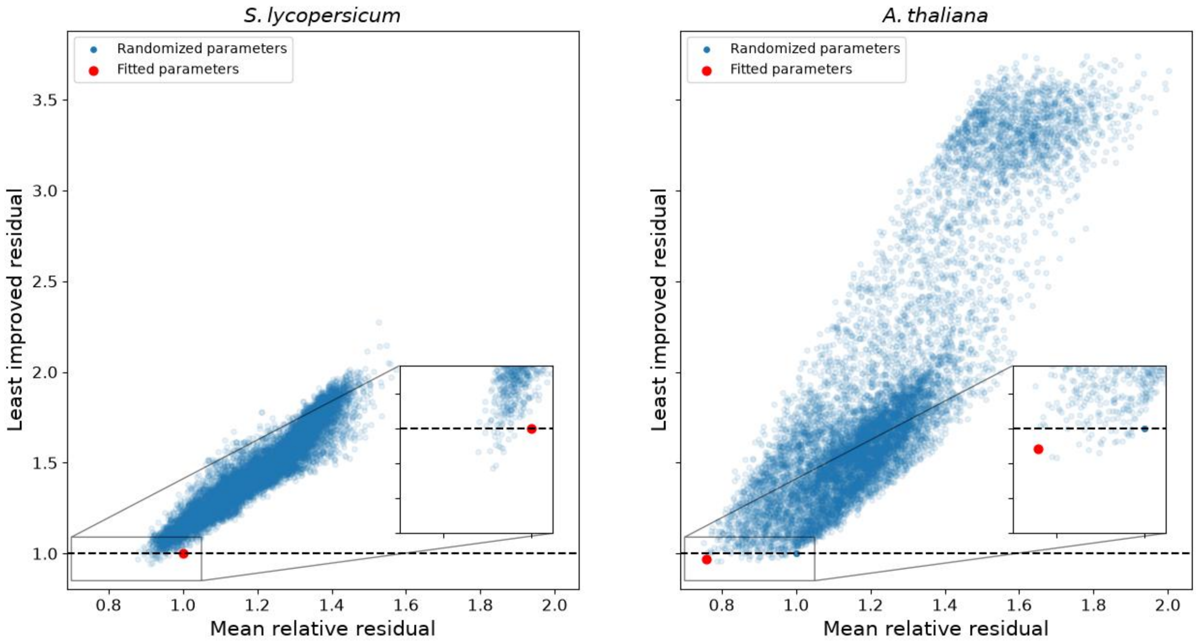


**Fig. S15 Monte Carlo simulation on selected parameter sets.** For each fitted parameter set of *A. thaliana* (left) and tomato (right), ~10,000 parameters sets were randomly sampled within ±10% of the fitted parameter values. Each randomized parameter set was used to run the model and respective residual errors were calculated from the differences between the model output and the measured data of NPQ and quantum yield of photosystem II ($\Phi_{PSII}$). The plots show the normalized, maximum residual error relative to the residual error of the corresponding fitted parameter set (y-axis) against the normalized mean residual error calculated from the same randomized parameter set (x-axis). In both cases, only a small subset of randomized parameter sets performed better than the fitted parameter sets with normalized residual errors ≤1. This indicates that fitted parameter sets represent locally robust solutions and that the observed fit is not readily obtained by small random perturbations of the fitted parameters. One of those better performing sets was chosen for further analysis of *A. thaliana*, because it further improved the fit. For tomato the initial parameter set was kept, because none of the tested parameter sets substantially improved the fit.

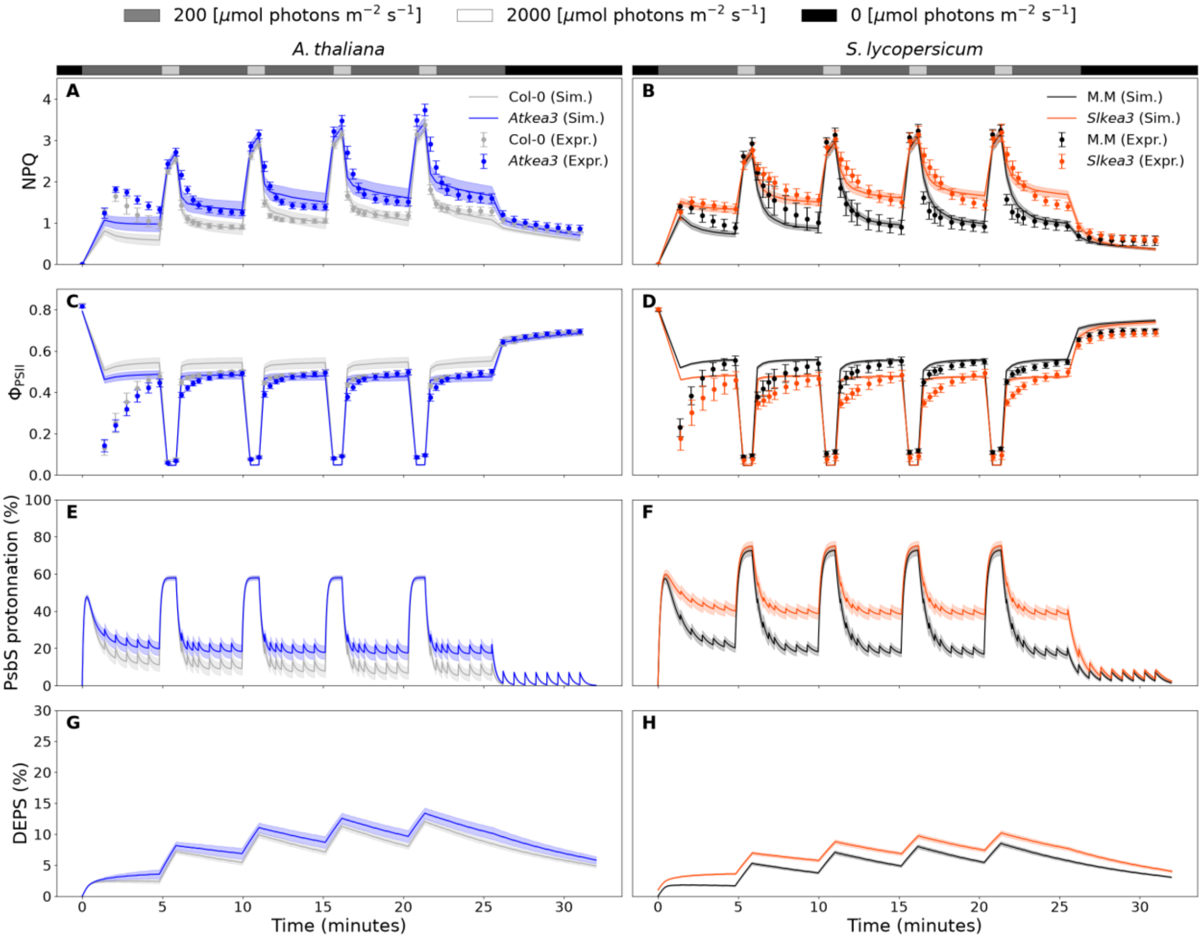


**Fig. S16 Modelled dynamics of selected photosynthetic parameters during light intensity fluctuations.** The full simulated results of NPQ (A-B), photosystem II quantum efficiency ($\Phi_{PSII}$, C-D), PsbS protonation state (E-F) and xanthophyll cycle de-epoxidation state (DEPS, G-H) of WT (Col-0 or M.M.) and *kea3* mutant of *A. thaliana* and tomato (*S. lycopersicum*) over four light intensity fluctuations of 4 min low light (200 µmol photons $m^{-2}$ $s^{-1}$) and 1 min high light (2000 µmol photons $m^{-2}$ $s^{-1}$), a final 4 min low light phase and 5 min darkness are shown.

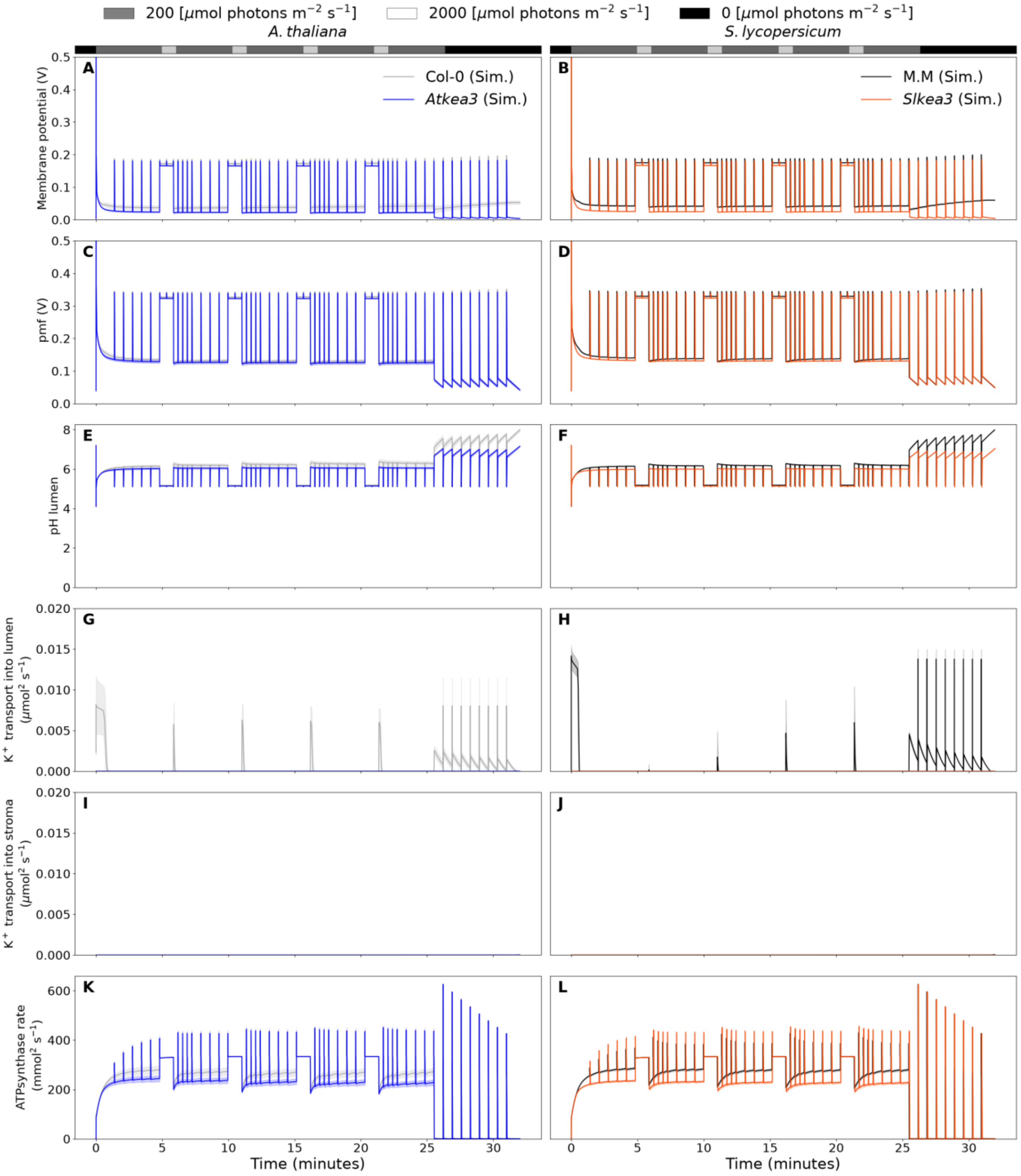


**Fig. S17 Modelled dynamics of selected KEA3-related parameters during light intensity fluctuations.** The full simulated results of membrane potential (A-B), PMF (C-D), lumen pH (E-F) KEA3-depemdemt K+ transport into the lumen (G-H), KEA3-dependent transport of K+ out of the lumen (I-J) and rate of the ATP synthase (K-L) of WT (Col-0 or M.M.) and *kea3* mutant of *A. thaliana* and tomato (*S. lycopersicum*) over four light intensity fluctuations of 4 min low light (200 µmol photons m$^{-2}$ s$^{-1}$) and 1 min high light (2000 µmol photons m$^{-2}$ s$^{-1}$), a final 4 min low light phase and 5 min darkness are shown.

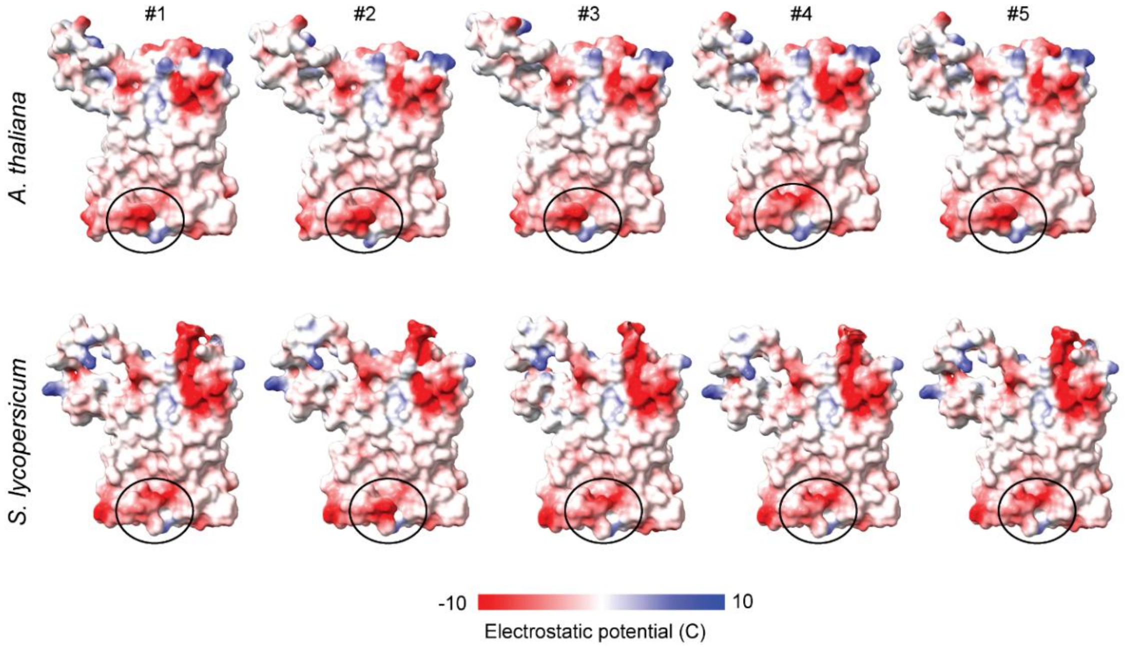


**Fig. S18 PsbS structure predictions and surface electrostatic potential.** Alphafold 3 was used to predict 5 different PsbS structures per species (*A. thaliana* and tomato*, S. lycopersicum*), which were visualized for surface electrostatic potential using ChimeraX. For each species, 4 out of 5 predictions take on different surface charge around the pH-responsive glutamate positioned between helices 3 and 4 (see Fig. 8).

### Methods S1 Construction of mathematical models

The ordinary differential equations (ODEs)-based mechanistic kinetics model contains 19 equations (Eq. 1-19), 13 state variables (Table S1), and 67 parameters (Table S2). It builds on the NPQ model of Matuszyńska et al. (Matuszyńska et al., 2016), which describes the qE and PSII activity. Linear electron flow (LEF) is represented in a simplified form without an explicit description of PSI. Our implementation further incorporates a description of proton motive force (*PMF)* from (Cruz et al., 2001) and KEA3 activity based on (Davis et al., 2017) and Li et al. (Li et al., 2021), with additional regulatory features of KEA3 based on experimental work (Uflewski et al., 2024).

**Model assumptions**

The model inherits several assumptions from the underlying models. Because the model is intended to describe short-term NPQ responses, protein abundances are assumed to remain constant and changes in gene expression are not considered. Stromal pH and the lumen buffering capacity are also assumed to remain constant. The chlorophyll concentration within the thylakoid membrane and the lumen-to-stroma volume ratios relative to thylakoid area were adopted from previous publications. These parameters may differ between *A. thaliana* (At) and *S. lycopersicum* (Sl) and were therefore treated as fixed model parameters. The xanthophyll pool and PsbS protein content are normalized, such that their modelled values represent relative abundances within their respective total pools rather than absolute concentrations.

**Updated modules**

*PMF* description

The membrane potential (Eq. 8) is calculated based on the differential equation:

$$\frac{dV}{dt} = \frac{1}{C} \cdot \frac{dQ}{dt} = \frac{z \cdot F}{C} \cdot \frac{dJ}{dt}$$

with $V$ membrane potential, $C$ membrane capacitance, $F$ Faraday constant, $z$ ion valence, $J$ ion fluxes.

Photosystem II

We keep the 4-state description from the previous model (Matuszyńska et al., 2016), However, instead of the quasi-state assumption for PSII, which assumes the transitions between the 4 states are significantly faster than the chemical reaction, in this model, we describe the states of PSII dynamically (Eq 1-3).

Cytochrome $b_6f$ activity

We keep the previous equation; however, the equilibrium constant is calculated based on *PMF* instead of the pH difference, and the final rate is constrained via pH regulation. For the pH regulation, we implemented the equation from (Cruz et al., 2001).

$$pHmod = 1 - \frac{1}{10^{pH_{lumen} - pKreg} + 1}$$

ATP synthase activity

The fraction of active ATP synthase upon *PMF* activation, is calculated as:

$$ATPsynthase_{activity,pmf} = \frac{ATPsynthase_{active}}{ATPsynthase_{total}} = \frac{e^x}{1 + e^x}$$

$pKE_0$ and $b$ are the fitted parameters to present the activation effect of *PMF* on ATP synthase activity (Eqs. 28)(Junesch and Gräber, 1987).

NPQ components

In this model, we only model the qE component of NPQ, as it is the major component of NPQ recorded via PAM measurement. We keep the same 4-state assumption about qE from the previous model, but the equations are changed - all components now contribute to NPQ via the quencher activity, described by first-order mass action kinetics. This is in agreement with the current assumption of the NPQ component.

$$Q = \gamma_0 \cdot Vx \cdot PsbS + \gamma_1 \cdot Vx \cdot PsbS^P + \gamma_2 \cdot Zx \cdot PsbS + \gamma_3 \cdot Zx \cdot PsbS^P$$

KEA3 activity

For the representation of KEA3 in our model, we assumed that transport can occur in either direction and that the transport rate is the same in both directions. The stoichiometry of the forward and reverse reactions is adjusted based on the volume of stroma and lumen.Low pH in the lumen inhibits KEA3 activity, while high energy level in chloroplasts, represented by ATP concentration,- inhibits KEA3 activity. Stromal pH value and proton concentration are kept constant. The initial K+ concentration is set to 0.1 M in both lumen and stroma.

The rate of KEA3 is calculated via Eqs 35,36.

KEA3 activity regulation

KEA3 regulation via pH and ATP regulation is defined by:

$$KEA3_{activity} = KEA3_{activity(pHregulation)} \cdot KEA3_{activity(ATPregulation)}$$

$$KEA3_{activity(pHregulation)} = \frac{10^{(pH_{lumen} - pK_{KEA3})}}{(10^{((pH_{lumen} - pK_{KEA3})} + 1)}$$

with $pK_{KEA3}$ the assumed pKa value for KEA3. In previous publications KEA3 is shown to be potentially influenced by stroma pH. However, as the model keep a constant pH in the stroma, we used the pH of the lumen as an proxy for the pH regulation instead.

$$KEA3_{activity(ATPregulation)} = \frac{(1-c)}{(1 + e^{((ATP - ATP_{thres})/c)})}$$

with $ATP_{thres}$ representing the ATP level at which KEA3 activity is inhibited and $c$ to define the threshold level.

Table S1. Variables in the model

| Names | Description |
|---|---|
| State variables | |
| $B0$ | The ground state of PSII |
| $B1$ | The excited state of PSII with oxidized QA |
| $B2$ | The charge separation state of PSII with oxidized QA and not excited PSII |
| $PQH_2$ | $PQH_2$ level |
| $ATPase^*$ | Activity of ATPsynthase dependent on time |
| $ATP$ | ATP content |
| $H^+_{lumen}$ | Lumen proton concentration |
| $\Delta\psi$ | Membrane potential |
| $[K^+_{lumen}]$ | Lumen potassium ion concentration |
| $[K^+_{stroma}]$ | Stroma potassium ion concentration |
| $PsbS$ | Relative level of non-protonated PsbS protein |
| $Vx$ | Relative level of violaxanthin |
| Derived variables | |
| B3 | The excited state of PSII with oxidized QA |
| | PQ level |
| ADP | ADP content |
| $pH_{lumen}$ | Lumenal pH |
| $\Delta pH$ | pH difference between the lumen and the stroma |
| PMF | Proton motive force |
| $PsbS^P$ | Relative level of protonated PsbS protein |
| $Zx$ | Relative level of zeaxanthin |

**Differential equations**

[1] $\frac{dB0}{dt} = v_{B10} - v_{B01} + v_{B20}$

[2] $\frac{dB1}{dt} = v_{B01} - v_{B10} - v_{B12}$

[3] $\frac{dB2}{dt} = v_{B12} + v_{B32} - v_{B23} - v_{B20}$

[4] $\frac{dPQH_2}{dt} = v_{PSII} - v_{PQ_{ox}}$

[5] $\frac{dATPase^*}{dt} = v_{actATPase} - v_{deactATPase}$

[6] $\frac{dATP}{dt} = v_{ATPsynthase} - v_{ATP_{consumption}}$

[7] $b_H \cdot \frac{d[H^+_{lumen}]}{dt} = 2v_{B12} + 4v_{PQ_{ox}} - \frac{14}{3} v_{ATP_{synthase}} - v_{KEA3(forward)} - v_{leak}$

[8] $\frac{d\Delta\psi}{dt} = Volts\_per\_charge \cdot (\frac{4}{b_H} v_{PQ_{ox}} + \frac{2}{b_H} v_{B12} - \frac{HPR}{b_H} v_{ATPsynthase} - \frac{1}{b_H} v_{leak})$

[9] $\frac{d[K^+_{lumen}]}{dt} = v_{KEA3(forward)} - v_{KEA3(reverse)}$

[10] $\frac{d[K^+_{stroma}]}{dt} = v_{KEA3(reverse)} - v_{KEA3(forward)}$

[11] $\frac{dPsbS}{dt} = v_{PsbS} - v_{PsbS^P}$

[12] $\frac{dVx}{dt} = -v_{X_{Epox}} + v_{X_{Deepox}}$

Rate equations

[13] $v_{actATPase} = k_{actATPase} \cdot H(PFD) - k_{deactATPase} \cdot (1 - H(PFD)) \cdot ATP_{ase}^*$

[14] $v_{B12} = k_P \cdot 0.5 \cdot B1$

[15] $v_{B10} = (k_F + (k_{H_{Qslope}} \cdot Q + k_{H0})) \cdot B1$

with

[16] $Q = \gamma_0 \cdot Vx \cdot PsbS + \gamma_1 \cdot Vx \cdot PsbS^P + \gamma_2 \cdot Zx \cdot PsbS + \gamma_3 \cdot Zx \cdot PsbS^P$

[17] $v_{B01} = PPFD \cdot B0$

[18] $v_{B23} = PPFD \cdot B2$

[19] $v_{B20} = k_{PQred} \cdot PQ \cdot B2 - \frac{k_{PQred}}{K_{eq,PQH2}} \cdot PQH2 \cdot B0$

with

[20] $K_{eq,PQH2(pH)} = e^{\frac{-(2 \cdot F \cdot E^0(QA/QA^-) - 2 \cdot F \cdot E^0(PQ/PQH_2) + 2 \cdot log(10) \cdot RT \cdot pH_{stroma})}{RT}}$

[21] $v_{PQ_{ox}} = v_{b6f} + v_{PTOX}$

$$(PPFD * k_{b6f} * pHmod)(\frac{K_{eq,PQH2}}{K_{eq,PQH2} + 1} \cdot PQH2 - \frac{1}{K_{eq,PQH2} + 1} \cdot PQ) + k_{PTOX} \cdot O_{2,ex} \cdot PQH2$$

with

[22] $pHmod = 1 - \frac{1}{10^{pH_{lumen} - pKreg} + 1}$

[23] $Keqcytb6f = e^{\frac{\Delta G_{PQox}}{RT}}$

for

[24] $\Delta G_{PQox} = 2\Delta G_{PQ,PQH2} + \Delta G_{PCP700} + \Delta G_{H+}$

$-(-2 \cdot F \cdot E^0_{PQ/PQH_2} + 2 \cdot log10 \cdot RT \cdot pH_{lumen}) + (-F \cdot E^0_{PQ/PQH_2}) + 2 \cdot F \cdot pmf$

[25] $v_{ATPsynthase} = ATP^*_{ase} \cdot ATPsynthase_{activity,pmf} \cdot k_{ATPsynthase} \cdot (ADP - \frac{ATP}{K_{eq,ATPsynthase}})$

with

[26] $K_{eq,ATPsynthase} = Pi_{mol} * e^{\frac{-\Delta G_{ATP}}{RT}}$

[27] $\Delta G_{ATP} = \Delta G^0_{ATP} - HPR \cdot pmf \cdot F$

The fraction of active ATP synthase is calculated:

[28] $ATPsynthase_{activity,pmf} = \frac{ATPsynthase_{active}}{ATPsynthase_{total}} = \frac{e^x}{1+e^x}$

with

[29] $x = log(10^{-pKE_0}) + b \cdot \frac{\tilde{\mu}_{H+}}{RT}$

$\tilde{\mu}_{H+}$ is the proton electrochemical potential across the thylakoid membrane.

[30] $\tilde{\mu}_{H+pmf \cdot F}$

[31] $pmf = \Delta\psi + ln(10) \cdot \frac{RT}{F} \cdot \Delta_{pH}$

[32] $\Delta_{pH} = pH_{stroma} - pH_{lumen}$

[33] $v_{leak} = k_{leak} \cdot ([H^+]_{stroma} - [H^+])$

[34] $v_{Prot} = k_{Prot} \cdot \frac{[H_{lumen}]^{nHL}}{[H_{lumen}]^{nHL} + pH_{SatLHC}{}^{nHL}} \cdot PsbS$

[35] $v_{Deprot} = k_{Deprot} \cdot \frac{Zx^{nHX}}{Zx^{nHX} + K_{Zsat}{}^{nHX}} \cdot PsbS^P$

[36] $v_{X_{Epox}} = k_{EpoxZ} \cdot Zx$

[37] $v_{X_{Deepox}} = k_{DeepoxV} \cdot \frac{[H_{lumen}]^{nHX}}{[H_{lumen}]^{nHX} + pH_{SatZ}{}^{nHX}} \cdot Vx$

[38] $v_{KEA3(forward)} = \frac{V_{lumen}}{V_{stroma}} \cdot k_{KEA3} \cdot KEA3_{activity} \cdot ([K^+_{stroma}] \cdot [H^+_{lumen}] - [K^+_{lumen}] \cdot [H^+_{stroma}])$

[39] $v_{KEA3(reverse)} = \frac{V_{stroma}}{V_{lumen}} \cdot k_{KEA3} \cdot KEA3_{activity} \cdot ([K^+_{lumen}] \cdot [H^+_{stroma}] - [K^+_{stroma}] \cdot [H^+_{lumen}])$

KEA3 regulation by pH and ATP is calculated:

[40] $$KEA3_{activity} = KEA3_{activity(pHregulation)} \cdot KEA3_{activity(ATPregulation)}$$

[41] $$KEA3_{activity(pHregulation)} = \frac{10^{(pH_{lumen} - pK_{KEA3})}}{(10^{((pH_{lumen} - pK_{KEA3})} + 1)}$$

with $pK_{KEA3}$ the assumed pKa value for KEA3.

[42] $$KEA3_{activity(ATPregulation)} = \frac{(1-c)}{(1+e^{((ATP - ATP_{thres})/c)})}$$

<u>Other derived quantities</u>

We derive the equation to calculate the fluorescence signal from PSII:

[43] $$Fluorescence = \frac{k_F}{(k_{H_{Qslope}} \cdot Q + k_{H0}) + k_F + k_P} \cdot B_0 + \frac{k_F}{(k_{H_{Qslope}} \cdot Q + k_{H0}) \cdot Q + k_F} \cdot B_2$$

We calculated the changes in membrane potential caused by a single charge movement via the amount of chlorophyll per area of thylakoid membrane, the Faraday constant and the thylakoid membrane specific capacitance $C$ (Li et al., 2021):

[44] $$Volts\_per\_charge = \frac{[Chl](mol \cdot m^{-2}) \cdot F}{C(F \cdot m^{-2})}$$

The proton concentration in the model is calculated based on the amount of protons and the respective volume:

[45] $$H^{+}_{lumen/stroma}(M) = \frac{[H^{+}_{lumen/stroma}](mmol \cdot molChl^{-1})/1000}{lumen/stromavolumnperareamembrane(L \cdot m^{-2})} \cdot [Chl](mol \cdot m^{-2})$$

S2. Parameters used in the model

| Parameters | | | | | |
|---|---|---|---|---|---|
| Parameter | Value | Unit | Description | Species of origin | Ref. |
| Pool sizes | | | | | |
| $PSII^{tot}$ | 2.5 | $mmol \cdot (molChl)^{-1}$ | Total pool of PSII reaction centers | Tobacco (*Nicotiana tabacum*) | (Schöttler et al., 2004) |
| $PQ^{tot}$ | 20 | $mmol \cdot (molChl)^{-1}$ | Total plastoquinone pool $(PQ + PQH_2)$ | Spinach (*Spinacia oleraceae*) | (Kirchhoff et al., 2002) |
| $AP_{tot}$ | 50 | $mmol \cdot (molChl)^{-1}$ | total adenosine phosphate pool (ATP + ADP) | Spinach (*Spinacia oleraceae*) | (Heineke et al., 1991) |
| $PsbS_{tot}$ | 1 | dimensionless | Relative abundance. PsbS normalized, under the assumption that the amount of PsbS remains constant over the simulation time | | (Matuszyńska et al., 2016) |
| $X_{tot}$ | 1 | dimensionless | Total pool of xanthophyll (Vx + Ax + Zx) normalized, under the assumption that the total amount of xanthophyll remains unchanged | | (Matuszyńska et al., 2016) |
| $O_2^{tot}$ | 8 | $mmol \cdot (molChl)^{-1}$ | external oxygen pool, corresponds to 250 $\mu$mol $L^{-1}$ | | (Ebenhöh et al., 2014) |
| $Pi_{mol}$ | 0.01 | $mmol \cdot (molChl)^{-1}$ | internal pool of phosphate, required to calculate ATP equilibrium | | (Matuszyńska et al., 2016) |
| Rate constants and key parameters | | | | | |
| Photosystem II | | | | | |
| $k_{H0}$ | $5 \cdot 10^8$ | $s^{-1}$ | rate of basal non-radiative decay | Estimated | |
| $k_{H_{Qslope}}$ | $5 \cdot 10^9$ | $s^{-1}$ | coefficient to calculate heat dissipation rate from quencher activity | Estimated | |
| $k_F$ | $6.25 \cdot 10^8$ | $s^{-1}$ | rate of fluorescence | | (Ebenhöh et al., 2014) |
| $k_P$ | $6.94 \cdot 10^9$ (Sl)<br>$6.35 \cdot 10^9$ (At) | $s^{-1}$ | rate of photochemistry | Fitted | |
| Cytochrome b6f and plastoquinone pool kinetics | | | | | |

| | | | | | |
|---|---|---|---|---|---|
| $k_{Cytb6f}$ | $0.022$ (Sl)<br>$0.205$ (At) | $mmol^{-2} \cdot (molChl)^2 s^{-1}$ | rate constant representing all oxidation reaction starting from Cytb6f, fitted with experimental data | Fitted | (Matuszyńska et al., 2016) |
| $pKreg$ | $6.3$ | dimensionless | pKa for the PCON process when Cytb6f get down regulated via low lumenal pH | | (Cruz et al., 2001) |
| $k_{PQ_{red}}$ | $250$ | $mmol \cdot (molChl)s^{-1}$ | estimated value to ensure the half-life from QB→PQ ≈ $1ms$ | Spinach (*Spinacia oleraceae*) | (De Wijn and Van Gorkom, 2001;<br>Ebenhöh et al., 2014) |
| $k_{PTOX}$ | $0.01$ | $mmol^{-1} \cdot (molChl)s^{-1}$ | rate constant of PTOX | *Chlamydo monas sp.* | (Ebenhöh et al., 2014) |
| ATPsynthase | | | | | |
| $k_{ActATPase}$ | $0.01$ | $s^{-1}$ | rate constant of activation of the ATPase in the light | | (Matuszyńska et al., 2016) |
| $k_{DeactAtpase}$ | $0.002$ | $s^{-1}$ | parameter relating the deactivation of the ATPase at night | | (Matuszyńska et al., 2016) |
| $pKE0$ | $7.21$ (Sl)<br>$6.61$ (At) | dimensionless | parameter for *PMF*-related ATP synthase activation | Fitted | (Junesch and Gräber, 1987) |
| $b$ | $3.1925$ (Sl)<br>$3.029$ (At) | dimensionless | parameter for *PMF*-related ATP synthase activation | Fitted | (Junesch and Gräber, 1987) |
| $k_{ATPsynthase}$ | $20$ | $s^{-1}$ | "fitted to allow a maximal production of ≈ $400 mmol(molChl)^{-1}s^{-1}$, which allows for the maximal reported CO2 fixation rate of 500 $\mu mol$ reduced CO2$(\mu molChl)^{-1}h^{-1}$" | Spinach (*Spinacia oleraceae*) | (Dietz and Heber, 1984;<br>Ebenhöh et al., 2014) |
| $HPR$ | $14/3$ | dimensionless | ratio of protons to ATP in ATP synthase | | (Nicholls and Ferguson, 2002) |
| $\Delta G_{0_{ATP}}$ | $30.6$ | $kJ/mol/RT$ | standard Gibbs free energy change of ATP formation | | (Nicholls and Ferguson, 2002) |
| Other processes | | | | | |
| $k_{leak}$ | $1000$ | $s^{-1}$ | rate constant of the proton flow from chloroplast thylakoid lumen to stroma | | (Matuszyńska et al., 2016) |
| $k_{ATPconsumptio}$ | $10s^{-1}$ | $s^{-1}$ | Rate of ATP consumption | | (Ebenhöh et al., 2014) |
| Parameters associated with the quencher activity | | | | | |
| Parameters associated with xanthophyll cycle | | | | | |
| $k_{DeepoxV}$ | $0.000672$ (Sl)<br>$0.000905$ (At) | $s^{-1}$ | rate of Vx deepoxiation | Fitted | |

| | | | | | |
|---|---|---|---|---|---|
| $k_{EpoxZ}$ | 0.0018664 (Sl) 0.001614 (At) | $s^{-1}$ | rate of Zx epoxidation | Fitted | |
| $K_{pHSat}$ | 5.8 | dimensionless | half-saturation pH for de-epoxidase activity, highest activity at pH 5.8 | | (Matuszyńska et al., 2016) |
| $nH_X$ | 5 | dimensionless | Hill-coefficient for the protonation kinetics of VDE | | (Matuszyńska et al., 2016) |
| $nH_Z$ | 3 | dimensionless | Hill-coefficient for the inhibition effect of Zx on PsbS deprotonation process | Fitted | |
| $K_{ZSat}$ | 0.35 (Sl) 0.585 (At) | dimensionless | half-saturation constant for the inhibition effect of Zx on PsbS deprotonation process | Fitted | |
| Parameters associated with PsbS protonation | | | | | |
| $nH_L$ | 3 | dimensionless | Hill-coefficient for activity of protonation | | (Matuszyńska et al., 2016) |
| $k_{ProtonationL}$ | 0.07392 (Sl) 0.09938 (At) | $s^{-1}$ | rate of PsbS protonation | Fitted | |
| $k_{Deprotonation}$ | 0.02688 (Sl) 0.07103(At) | $s^{-1}$ | rate of PsbS de-protonation | Fitted | |
| $K_{phSatLHC}$ | 6.3 | dimensionless | pKa of PsbS activation | | (Zaks et al., 2012) |
| Parameters associated with KEA3 activity | | | | | |
| $k_{KEA3}$ | 28 (Sl) 29 (At) | $s^{-1}$ | influence of protein KEA3 to the proton efflux from thylakoid lumen | Fitted | |
| $K_{lumen\_conc}$ | 0.01 | $M$ | Initial lumen potassium concentration | | (Li et al., 2021) |
| $K_{stroma\_conc}$ | 0.01 | $M$ | Initial stroma potassium concentration | | (Li et al., 2021) |
| $pK_{KEA3}$ | 6.75 (Sl) 7.07 (At) | dimensionless | the pKa for pH regulation of KEA3 activity | Fitted | |
| $ATP_{thres\_KEA3}$ | 20.5 (Sl) 21.89 (At) | $mmol.(molChl)s^{-}$ | Level of ATP that KEA3 start getting inhibited | Fitted | |
| Fitted quencher contribution factors | | | | | |
| $\gamma_0$ | 0.1 (Sl) 1.06 (At) | dimensionless | corresponds to base quenching not associated with protonation or zeaxanthin | Fitted | |
| $\gamma_1$ | 9.5 (Sl) 9.74 (At) | dimensionless | fast quenching present due to protonation | Fitted | |
| $\gamma_2$ | 6.0 (Sl) | dimensionless | fastest possible quenching | Fitted | |

| | | | | | |
|---|---|---|---|---|---|
| | 6.18 (At) | | | | |
| $\gamma_3$ | 2.5 (Sl)<br>3.37 (At) | dimensionless | slow quenching of Zx present despite lack of protonation | Fitted | |
| Physical constants | | | | | |
| $F$ | 96.485 | $kJ$ | Faraday constant | | |
| $R$ | 0.0083 | $kJK^{-1}mol^{-1}$ | universal gas constant | | |
| $T$ | 298 | $K$ | temperature | | |
| $e$ | 2.71828 | dimensionless | | | |
| $PPFD$ | $Adjusted$ | $\mu mol\ photons\ s^{-1}\ m^{-2}$ | Light intensity | | |
| Standard potentials | | | | | |
| $E^0$ $(QA/QA^-)$ | $-0.140$ | $V$ | Standard redox potential of $QA/QA^-$ | Spinach (*Spinacia oleraceae*) | (Suzuki et al., 1987) |
| $E^0$ $(PQ/PQH_2)$ | 0.354 | $V$ | Standard redox potential of $PQ/PQH_2$ | Spinach (*Spinacia oleraceae*) | (Allakhverdiev et al., 2011) |
| $E^0$ $(PC/PC^-)$ | 0.380 | $V$ | Standard redox potential of $PC/PC^-$ | *Egeria densa* | (Okayama, 1976) |
| Chloroplast parameters | | | | | |
| $pH_{stroma}$ | 7.8 | dimensionless | Stromal pH | | (Cruz et al., 2001;<br>Matuszyńska et al., 2016) |
| $b_H$ | 100 | dimensionless | Lumen buffering capacity | | (Cruz et al., 2001;<br>Laisk et al., 2007) |
| Lumen volume | 0.0014 | $L \cdot m^{-2}$ | Lumen volume | | (Laisk et al., 2007) |
| Stroma volume | 0.0112 | $L \cdot m^{-2}$ | Stroma volume | | (Laisk et al., 2007) |
| Chl concentrati on | $350 \cdot 10^{-6}$ | $mol \cdot m^{-3}$ | Amount of chlorophyll per thylakoid membrane area | | (Laisk et al., 2007) |
| C | $0.6 \cdot 10^{-2}$ | $F \cdot m^{-2}$ | Thylakoid membrane capacitance | Calculated | (Davis et al., 2017;<br>Li et al., 2021) |
